# HOTARU: Automatic sorting system for large-scale calcium imaging data

**DOI:** 10.1101/2022.04.05.487077

**Authors:** Takashi Takekawa, Masanori Nomoto, Hirotaka Asai, Noriaki Ohkawa, Reiko Okubo-Suzuki, Khaled Ghandour, Masaaki Sato, Masamichi Ohkura, Junichi Nakai, Shin-ichi Muramatsu, Yasunori Hayashi, Kaoru Inokuchi, Tomoki Fukai

## Abstract

Currently, calcium imaging allows long-term recording of large-scale neuronal activity in diverse states. However, it remains difficult to extract neuronal dynamics from recorded imaging data. In this study, we propose an improved constrained nonnegative matrix factorization (CNMF)-based algorithm and an effective method to extract cell shapes with fewer false positives and false negatives through image processing. We also show that the evaluation metrics obtained during image and signal processing can be combined and used for false-positive cell determination. For the CNMF algorithm, we combined cell-by-cell regularization and baseline shrinkage estimation, which greatly improved its stability and robustness. We applied these methods to real data and confirmed their effectiveness. Our method is simpler and faster, detects more cells with lower firing rates and signal-to-noise ratios, and enhances the quality of the extracted cell signals. These advances can improve the standard of downstream analysis and contribute to progress in neuroscience.

## 1 Introduction

A major goal of neuroscience is to understand interactions within large populations of neurons, including network dynamics and emergent behaviors. This necessitates the recording of diverse activities within large populations of neurons. In recent years, genetic indicators have been widely recorded by microscopy, and advances in genetic optics have greatly improved the signal strength of calcium-binding protein indicators (Chen et al., 2013). Currently, neuronal activity can be imaged in vivo in a larger region of the brain for longer periods (Cotton et al., 2013; Ahrens et al., 2013; Prevedel et al., 2014; Diego Andilla and Hamprecht, 2013). In addition to two-photon microscopy, the development of miniaturized microscopes such as microendoscopes has greatly expanded the area that can be visualized, such as the deep brain of a freely moving mouse, and the obtained data have multiplied(Flusberg et al., 2008; Ghosh et al., 2011; Ziv and Ghosh, 2015). Therefore, an efficient and accurate method is required to detect the spatial positions and shapes of neurons, as well as their temporal firing patterns, from the obtained imaging data.

A method used to extract individual cells from calcium imaging data is region of interest (ROI) analysis, which first extracts an ROI and then extracts the temporal changes in calcium concentration within the ROI (Smith and Häusser, 2010; Pinto and Dan, 2015; Barbera et al., 2016; Klaus et al., 2017). However, this method struggles to effectively separate signals from neurons that overlap in spatial distribution. Additionally, there is no accurate and efficient method for obtaining ROIs for large datasets (Resendez et al., 2016). Another method is PCA / ICA analysis (Mukamel et al., 2009). Because this method is a linear separation method, it cannot accurately separate cells with regard to spatial overlap (Resendez et al., 2016). Consequently, a third method, which includes analysis of nonnegative matrix factorization (NMF) (Maruyama et al., 2014) and constrained NMF (CNMF) (Pnevmatikakis et al., 2013, 2016) has been proposed. CNMF analysis models the position and shape of cells (footprint), the relationship between spikes and calcium fluctuations, and the statistical nature of noise. Therefore, it is possible to deconvolve overlapping cells into pure activity without causing mutual noise contamination. However, algorithms that simply optimize the model are computationally expensive. The standard solution uses an iterative algorithm that alternately improves calcium fluctuations and cell shapes. But instabilities that degrade accuracy arise during this iteration. For this reason, various computational innovations have been proposed. Among them, CNMF-E is a widely used high-performance method that considers the effects of background fluctuations in addition to numerous computational innovations (Zhou et al., 2018).

One of the prominent limitations of CNMF-E, a competitive method fundamental to our proposed approach, lies in the initial selection of candidate cells. In CNMF-E, a temporal downsampling of the video sequence is performed to calculate the maximum values, standard deviations, and pixel correlations. The peaks in these statistics are then identified as candidate cell locations, and the candidates are greedily chosen to ensure that there is no spatial overlap.

However, there are significant shortcomings with this selection strategy. Determining an appropriate threshold parameter is challenging and has a profound influence on outcomes. Furthermore, the method’s reliance on temporal summarization poses the risk of overlooking neurons with low firing rates or those exhibiting weak signal intensities.

An additional aspect of CNMF-E that warrants improvement is the reliability of its iterative algorithm and the methodology for setting appropriate parameters. In CNMF-E, the sparsity of cellular shapes and spikes is achieved through an L1 regularization term. However, there is no clear criterion for determining the value of this regularization parameter, and the results can vary significantly depending on its setting.

Moreover, while the method defines the problem as an optimization of the objective function and attempts iterative improvements, there is a potential downside. Specifically, as cell shapes, spike time series, and local background conditions are sequentially optimized, the estimates for these variables may deteriorate rather than improve. Therefore, it is often more practical to stop the iterative process after a predetermined number of cycles instead of waiting for it to convergence. This termination point introduces another layer of subjectivity to the algorithm.

Another crucial aspect that warrants reconsideration in the CNMF-E is the role of the temporal and spatial baseline in critical spike event inference. The same calcium time series can signify relatively high-frequency, continuous spiking events or low-frequency events with a poor signal-to-noise ratio, depending on the established baseline (Takekawa et al., 2017). However, in CNMF-E, the mathematical definition of this baseline is similar to that of cellular activity, with the only differences being the constraints and regularization terms. This introduces an additional layer of arbitrariness, as the trade-off between the baseline estimation and cellular activity is heavily influenced by parameter choices.

In this paper, we propose HOTARU (a High-performance Optimizer to extract spike Timing And cell location from calcium imaging data via lineaR impUlse), a simple, automatic calcium imaging data analysis system that improves on the above two limitations. First, multiple parameters of cell size are set within a certain range, and an initial footprint is formed using a Laplacian Gaussian (LoG) filter and a watershed cuts algorithm (Cousty et al., 2008), which is effective for blob detection (Lindeberg, 2013). Additionally, candidate cells are detected independently in every frame of the video to prevent them from missing those with low firing rates. This means that a huge number of cell candidates are generated in the first step, from which a greedy method that takes advantage of the size of cells with variations efficiently reduces the number of candidate cells to the minimum necessary. There are no arbitrary parameters in the algorithm, and it operates automatically.

The HOTARU system update routine builds on the ideas of CNMF, but with several improvements to address its limitations. First, we incorporate a novel group-normalized L1 regularization. In traditional L1 used in the CNMF, regularization is proportional to the footprint or intensity of spike trains. However, the actual intention is to impose constraints related to shape and firing frequency. With group-normalized L1 regularization, regularization is proportional to variables normalized to a maximum value of 1. We also propose a method to efficiently perform such a complex conditional optimization using the proximal gradient method. Second, we address the issue of background processing. For the overall background, we separate the temporal and spatial structures and process them in the form of dependent variables while adding L2 regularization. These processes stabilize the estimation of the baseline, preventing computational breakdown, especially in large-scale data where the matrix eigenvalues become small. Third, we introduce a local background that fundamentally adopts the same idea as in the CNMF-E. However, while CNMF-E intricately designs conditions for local backgrounds and cell candidates and iteratively optimizes them, HOTARU simultaneously optimizes multiple proximal operators, assuming that the local background is not constrained in its temporal structure.

In HOTARU, the primary parameters that require tuning are those representing the maximum and minimum values of the cell size radius. These values can be determined based on the necessary pre-computed statistics, rather than being arbitrary. As an option, we suggest adjusting the regularization term for spike trains and switching between cells and local backgrounds based on cell shape stability and the sparsity of spike trains. However, these operations are often unnecessary in many datasets. In addition, the impact of parameter changes can be quantitatively assessed, providing clear guidance when parameter adjustments are required.

In the following sections, we provide a detailed description of the method and present examples of its application to three experimental datasets comprising real data.

## 2 Mathematical models

Videos that capture calcium fluctuations from the brain can be represented as a matrix *F* [*t, x*] ∈ℝ^*T ×X*^, where *X* denotes the number of pixels and *T* represents the number of frames. Initially, we assume that cellular activities or localized background activities, observed as calcium fluctuations and denoted by *k*, can be represented as the outer product of spatial components *a*_*k*_[*x*] and temporal components *v*_*k*_[*t*]. For *a*_*k*_[*x*], we assume a spatial local structure and impose a non-negativity constraint, further constraining that the maximum value of *a*_*k*_[*x*] is 1. This prevents the ambiguity arising in *a*_*k*_[*x*]*v*_*k*_[*t*] for the same activity and is also related to the introduction of a normalization term by the probability model discussed below. As a result, the intensity of activity for element *k* is represented by *v*_*k*_[*t*]. Since the video contains significant noise in addition to meaningful activities, we collectively denote this noise as *ε*[*t, x*].The matrix is represented by following equation:

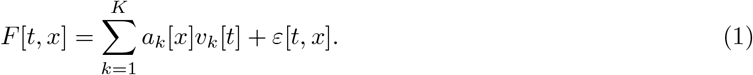

Up to this point, we have followed the basic structure of the CNMF. However, in this study, we treat the noise term *ε*[*t, x*] as an independent component without taking it as the outer product of temporal and spatial elements. Also, for cells and local background activities, while retaining the basic structure of the outer product, we alter the temporal and spatial constraints to efficiently perform calculations. We introduce distinct normalization terms for each component and optimize them simultaneously using the proximal gradient method. In CNMF-E, local background activity is treated separately from cells and is computed iteratively.

### 2.1 Celler and local background activities

Next, we discuss the component structures. We represent the calcium fluctuation that occurs when a cell produces a spike with the response function *γ*[*τ*]. We assume that the response to the spike train *u*_*k*_[*t*] is the linear sum of individual response functions. Furthermore, both the spike train and the response function are assumed to have non-negative values. In this paper, we used a double exponential function for the response function. However, the implementation can utilize any time-series response function. For the introduction of a subsequent regularization term, we constrain the maximum value of *u*_*k*_[*t*] to 1 and introduce *s*_*k*_ as the observed intensity for the entire cell. That is,

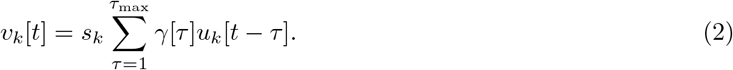

On the other hand, for local background activity, we do not make any assumptions about the temporal structure and do not impose a non-negativity condition. Constraints and normalization terms based on the probability models for both are described separately in the following sections.

### 2.2 Baseline regurization

We decompose distortions and fluctuations throughout the image, such as thermal noise, into temporal elements, spatial elements, and an overall baseline, denoted as

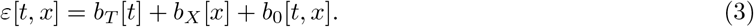

Specifically, we assume *b*_0_, *b*_*T*_ and *b*_*X*_ obays normal distributions:

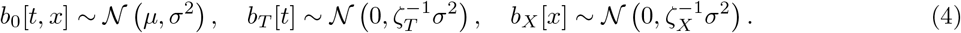

The baseline value, which is not directly evaluated, can be described based on the data and the activity of cells and other components under the principle of maximum likelihood estimation. The proportional relationship of the variances of *b*_0_, *b*_*t*_, and *b*_*x*_ based on parameters is practical and also contributes to the simplification of the evaluation function. However, it is important to note that due to the assumption of non-negativity of the cell components and the temporal constraints imposed by the response function, the structure is such that significant changes in activity estimation results can occur when the baseline *μ* changes. The constraints, such as regularization in calcium imaging models, can be rephrased as how to strike a balance between the intensity of cell activity and fluctuations in the baseline in a natural way.

### 2.3 Regularization by probabilistic models

The cellular shape *a*_*k*_[*x*] is locally present and becomes zero outside of this locality. This situation, although simplified, can be represented using an exponential probability model:

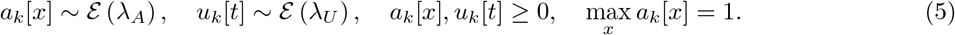

Assume that an exponential distribution corresponds to the regularization of L1 in a maximum a posteriori (MAP) estimation, resulting in a sparse solution. However, it is important to note that here, due to the constraint where max_*x*_ *a*_*k*_[*x*] equals 1, a creative solution approach to the solution is required. The assumption that max_*x*_ *a*_*k*_[*x*] is 1 signifies that this constraint does not depend on the intensity of the activity but is purely a constraint related to the “shape”. Similarly, for the spike sequence *u*_*k*_[*t*], since there is a standard firing frequency scale, it is assumed to follow an exponential distribution under the constraint max_*t*_ *u*_*k*_[*t*]. By employing this method, regularization can be performed in a normalized manner for each neuron candidate. Compared to evaluating *a*_*k*_[*x*]*u*_*k*_[*t*] as a whole using standard methods, this approach avoids the problem of becoming dependent on the intensity of activity. For the local background activity, we assume a Laplace distribution *ℒ* for *a*_*k*_[*x*]*v*_*k*_[*t*] as a whole:

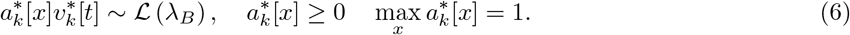

### 2.4 Fitting algorithm

In the following discussion, let *C* and *B* represent the sets of neuronal cells and the local background activity, respectively. If the *k*-th spatiotemporal element is estimated to be a neuronal cell, then *k ∈ C*. Similarly, if it is estimated to be part of the local background and is sparse, then *k ∈B*.

The MAP inference of model (Equation (1), (2)) and the prior (Equation (4), (5)) maximize the following logarithm of posterior likelihood *L*:

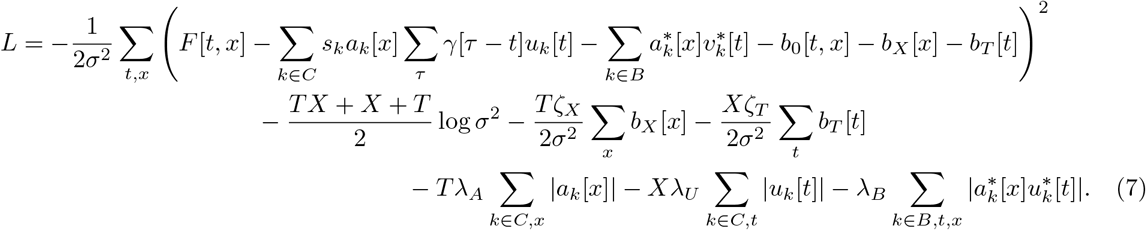

Here, we denote the parameters in a matrix form by *F ∈*ℝ^*X×T*^, 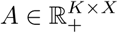, and *V ∈*ℝ^*K×T*^, respectively. Even if the data is centralized so that Σ_*x*_ *F* [*t, x*] = 0, Σ_*t*_ *F* [*t, x*] = 0, and Σ_*t,x*_ *F* [*t, x*]^2^ = *TX*, the generality of the solution is preserved. To maximize the likelihood function *L*, we determine the conditions for *b*_0_, *b*_*T*_, and *b*_*X*_. By performing variable elimination, *L* can be transformed into a simple form based on the error *E*:

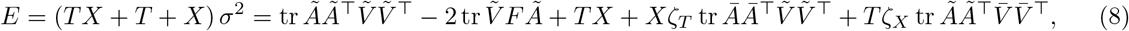

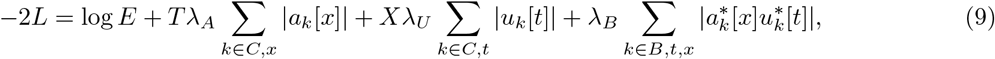

where *Ā* and 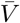 represent the average vector of *a*_*k*_ and *v*_*k*_, respectively. Similarly, 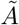 and 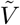represent the matrix of deviations obtained by subtracting the average from *a*_*k*_ and *v*_*k*_.

Then, by defining *â*_*k*_[*x*] = *s*_*k*_*a*_*k*_[*x*] and *û*_*k*_[*t*] = *s*_*k*_*u*_*k*_[*t*], we can split this problem into the following two sub-problems:

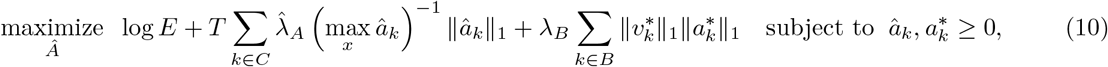

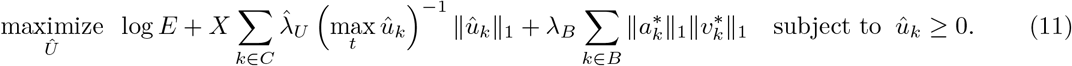

Both Equation (10) and (11) consist of a logarithm of quadratic form and a regularized L1 regularization term, which can be generalized as follows:

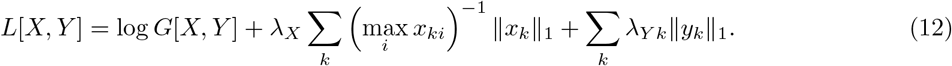

The proximal operator for this regularization term can be constructed as follows:

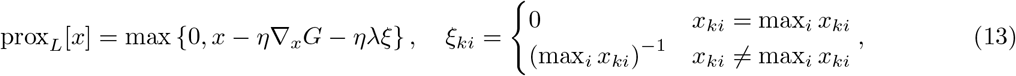

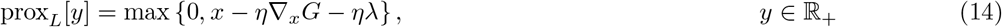

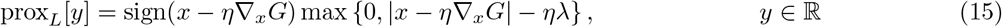

and the optimization problem can be solved by repeatedly applying the proximal operator. In addition, speed-ups, such as Nesterov acceleration, can be achieved during iterative application. In both Equation (10) and (11), the target function *g* is the quadratic form of the variable vector. Therefore, by precomputing the necessary matrices, only simple matrix products are needed in iteration. Consequently, the proposed algorithm can be processed at high speed using parallel operations of GPUs.

## 3 Image and signal processing

### 3.1 Peak finding by LoG filter

A Laplacian of Gaussian (LoG) filter, expressed by the following equation, is a spatial filter used in image processing to detect edges and blobs. This filter uses a size parameter *r*.

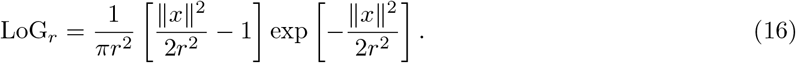

In our method, we use LoG filters with multiple size parameters 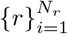 to create an initial footprint and shape, as well as evaluate the footprint obtained by an iterative method.

In creating the initial footprint, a filter denoted by 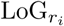 of different sizes *r*_*i*_ is applied to each frame *t* of the video *F* [*t*]:

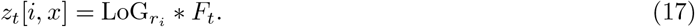

Then we collect all the local maxima (peaks) in (*i, x*). From the multitude of peaks identified by the greedy method, we select those peaks (*i, x*) that exhibit the highest ‘intensity’ *z*[*i, x*]. This selection is made such that the distance between any two chosen peaks is always less *θr*. The algorithm to efficiently obtain a set of peaks that satisfy this condition is described later. *θ* is a parameter that represents the distance between cells, otherwise expressed, the possible overlap of cells.

### 3.2 Segmentation of footprint

From the image filtered corresponding to the identified peak, we derive the initial footprint. We then apply the watershed method (Cousty et al., 2008) to the area surrounding the peak, enabling us to extract a region where a cell is present. For the iterative footprint, multiple LoG filters are similarly applied, the largest peak is selected, and the watershed method is used to calculate the area to be cut out. Each footprint *a*_*k*_ is limited to a maximum value of 1, thus the *z* value obtained by the LoG filters is interpreted differently than in the case of the initial footprint and is referred to as ‘firmness’ 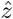 in our paper. The values obtained 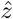 and *r* in the peak are useful for determining false positives.

Full details on the algorithms for initializing and then solving these three subproblems are provided in the Materials and methods section.

### 3.3 Evaluation Metrics for Results

In this study, we evaluate the integrity of the obtained cellular shapes (footprints) and spike time series using the following metrics. First, the maximum intensity *s* of the total cellular activity is considered. A higher value for *s* means that high fluorescence intensity is observed when spikes occur. Second, as a measure of cellular shape integrity, we introduce a metric called “firmness.” This is defined as the maximum value obtained when applying a Laplacian of Gaussian (LoG) filter to the footprint. While similar to the “intensity” used in initial cell detection, it differs in that it is calculated against a value *a* that is normalized to a maximum of 1. A firmness value ranging between 0.4 and 0.6 indicates that the cell shape is clearly estimated at the expected cell size, while lower values indicate distorted shapes or high levels of noise (see Figure 1 left).

**Figure 1:**
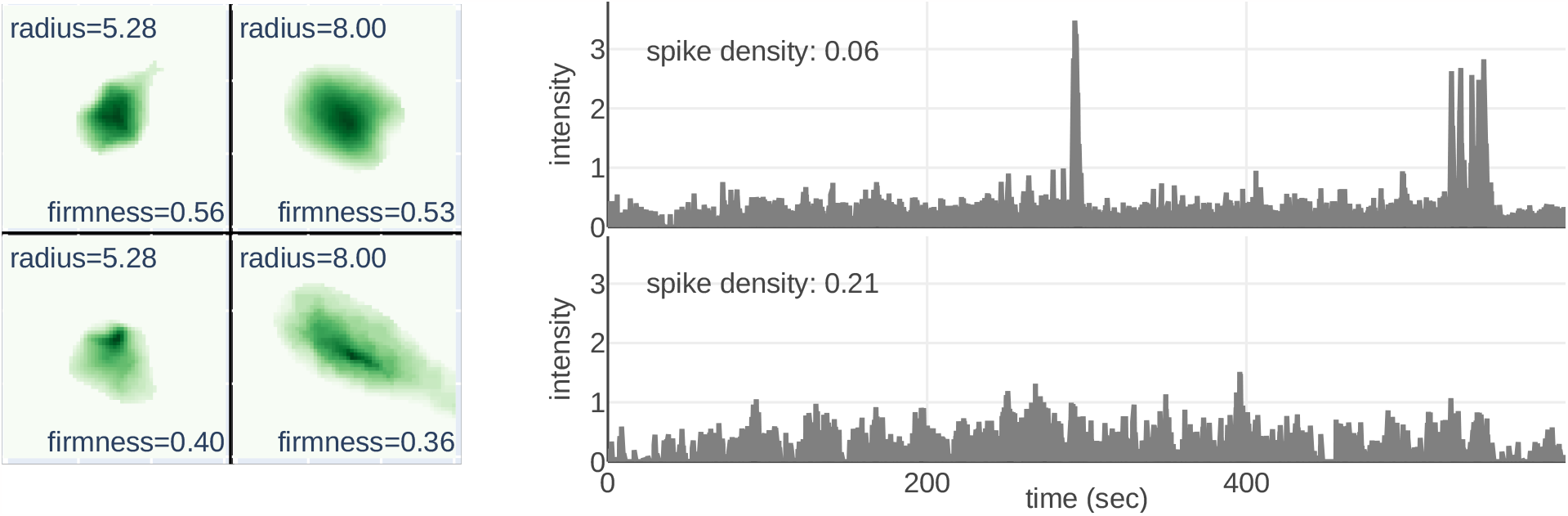
Illustrative examples of the evaluation metrics “footprint’s firmness” and “spike’s density.” Left: Showcases footprints with varying radii and firmness levels; higher firmness results in more distinct shapes. Right: Presents spike trains with different spike densities; density is generally inversely proportional to the signal-to-noise (S/N) ratio, and high density leads to indistinct spikes.

Lastly, the lack of integrity in the spike sequence is defined by the inverse of the signal-to-noise ratio (SNR). Considering that the signal is sparsely optimized, noise levels are robustly estimated from the time series *u* by using the median absolute deviation, with zero values excluded, to derive a standard deviation. In many cellular activities, this metric often falls below 0.2. However, in scenarios where signal intensity is weak or when activities have a longer time constant, such as vascular activities, this metric can have larger values (see Figure 1 right).

In the Results section, we demonstrate that the use of the proposed method leads to the selection of cell candidates with high signal intensity and footprint firmness while having low spike density. Importantly, candidates considered false positives are automatically excluded.

## 4 Results

In this paper, we apply a proposed analysis technique to three types of experimental data. We verify its accuracy and compare it to existing methods. First, we computed statistical metrics that summarize the basic properties of the data (see Figure 2). These metrics, including maximum value, standard deviation, and correlation with surroundings, are commonly used as potential indicators of cell activity in all frames of cell videos. Even from these summarized images, human observers can somewhat identify cells. Typically, peaks in specific metrics are sought as primary cell candidates, and a threshold is set to narrow the final cell candidate list. However, these methods often struggle to determine appropriate thresholds for noisy data. This can lead to issues such as “False Negatives,” where actual cells are missed, or “False Positives” where false signals are mistaken for cells.

**Figure 2:**
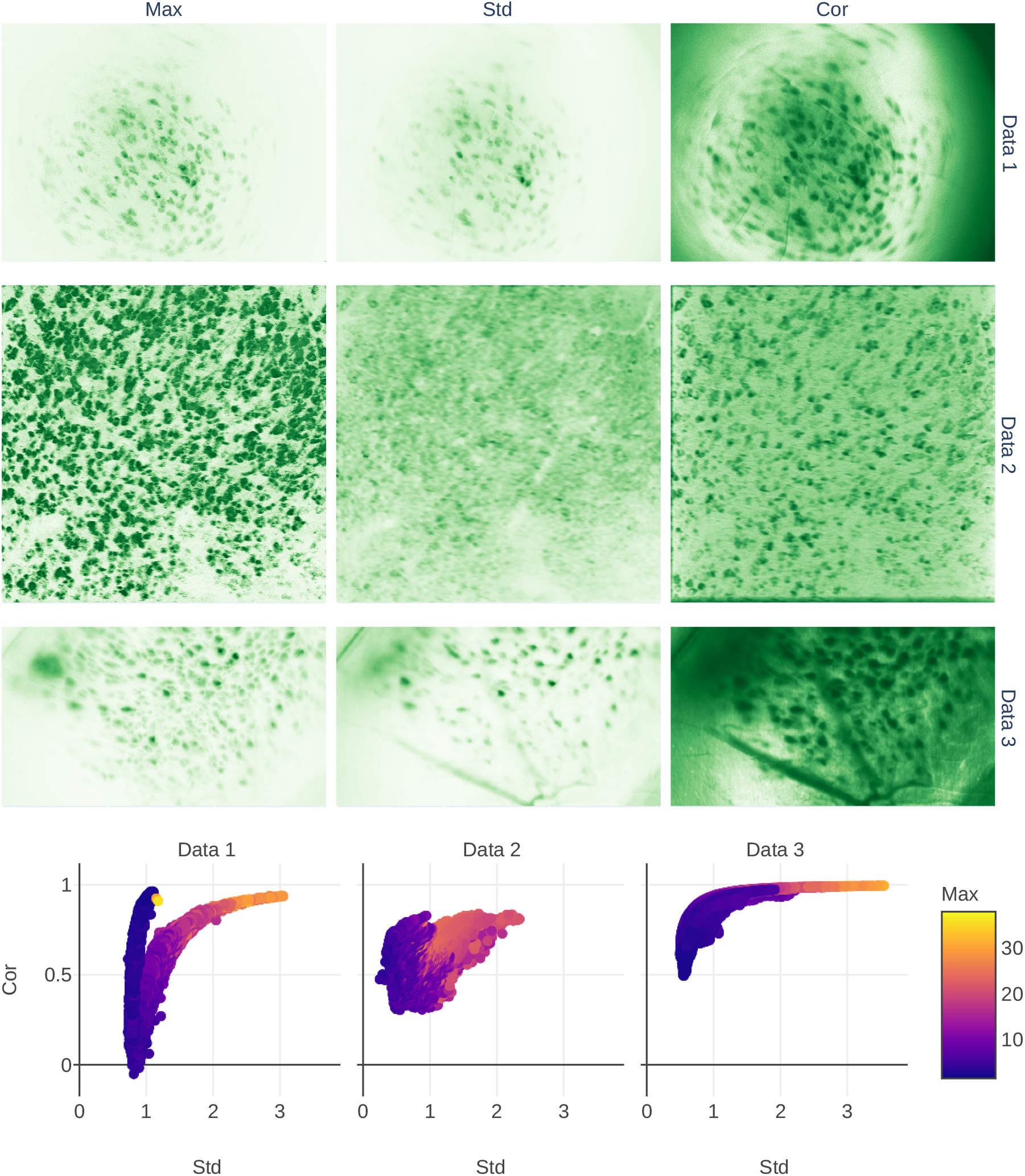
Simple statistical metrics derived from the actual experimental video data used in our analysis. The columns labeled “Max,” “Std,” and “Cor” represent the maximum pixel value, standard deviation of the pixel values, and correlation with neighboring pixels, respectively. Each row corresponds to Data 1, 2, and 3. Data 1 records CA1 using miniature microscopy; Data 2 captures the CA1 region of the dorsal hippocampus through two-photon microscopy; and Data 3 is a large-scale record of CA3 using miniature microscopy.

Data 1 is a dataset captured by miniature microendoscopy from the hippocampus of mice. It consists of 1,200 frames, each with dimensions of 720 *×* 540 pixels, recorded at 20 Hz over a span of 60 s. Data 2, on the other hand, are obtained by two-photon microscopy from the CA1 region of dorsal hippocampus of the mice. It encompasses 8,989 frames, each measuring 512*×* 512 pixels across an area of 532 *×* 532 μm, recorded at 15 Hz for 10 minutes (Sato et al., 2020). Data 3 is a larger set obtained by miniature microendoscopy from mice hippocampal CA3. This dataset comprises 74,000 frames of 607 *×* 344 pixel images, captured at 20 Hz, spanning 61 minutes.

Comprehensive experimental details for all three datasets can be found in the Materials Details section.

### 4.1 HOTARU reliably detect weak and low-firing-rate cells

To address the issues of false positives and negatives, we utilize the information from each frame rather than just summarized data. Using the LoG filter mentioned in the Methods section, we detect blobs of varying sizes that correspond to cells (see Figure 3). Applying the filter to all frames incurs computational costs, but by using a GPU, we can roughly process and apply the filter to about 1000 frames per second. This allows us to calculate the combination of frame t and radius r with the highest signal intensity for each pixel (x,y). Consequently, we can obtain a number of cell candidates equivalent to the image pixel count.

**Figure 3:**
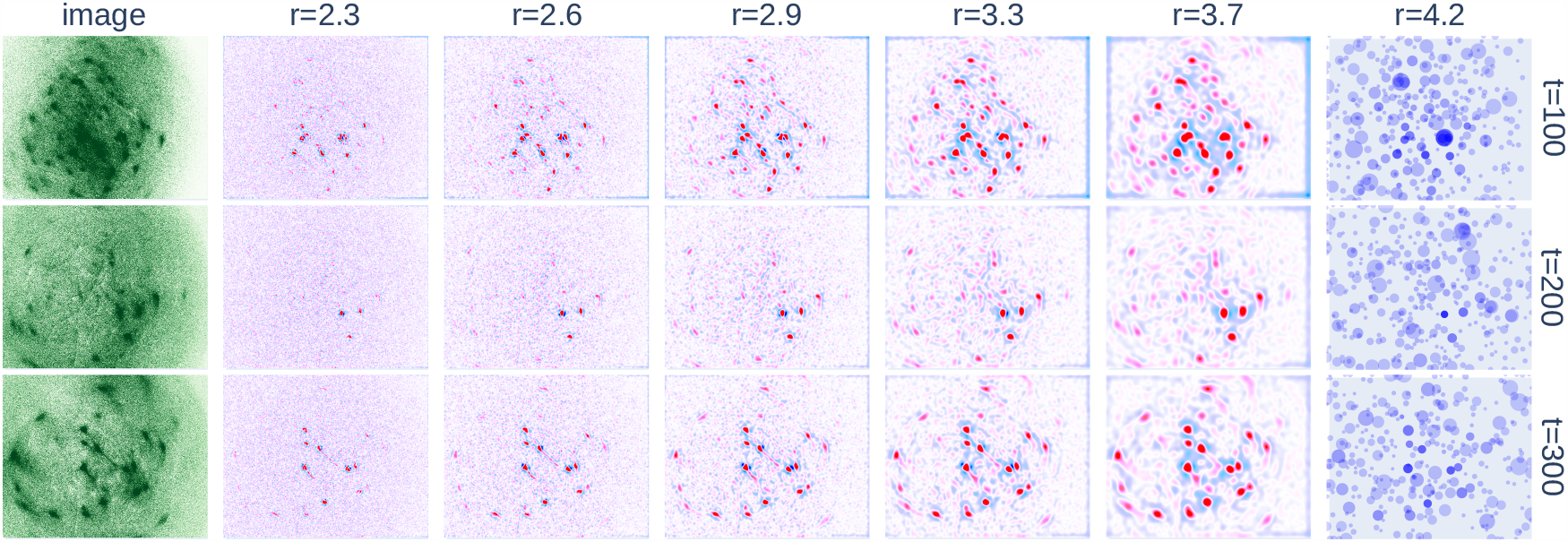
A summary of the results from applying multiple LoG (Laplacian of Gaussian) filters to each frame of Data 1. For frames at t=100, 200, and 300, the original images and results of applying LoG filters with radii r=2.3, 3.3, 4.8, 7.0, and 10.1 are displayed. The far-right column, labeled “peaks”, shows the detected maxima in a three-dimensional space across spatial dimensions (x,y) and radius r for each frame. The displayed circle’s radius corresponds to r, and its transparency corresponds to intensity.

Subsequently, the initial cell candidates are sorted based on their signal intensity. Candidates are filtered based on criterion that the circular region, which is proportional to its radius with a coefficient A, should not contain any other candidate with a higher signal intensity. Furthermore, not all the range of r from the initially applied LoG filter is adopted. We set a minimum *r*_min_ and a maximum *r*_max_. Candidates below *r*_min_ are ignored and those above *r*_max_ are treated as candidates for local background activity.

Considering the bidirectional conditions on the distance between cells, we can rigorously filter cell candidates that meet the conditions within a second using a greedy algorithm. Therefore, it is feasible to thoroughly adjust parameters such as *r*_min_, *r*_max_, and coefficient *θ* to ensure that the initial conditions are appropriate.

From Figure 4, the distribution of radius r and intensity indicates a high peak around the standard cell size of 4-10 pixels. By confirming this unimodal distribution for each dataset, setting *r*_min_ and *r*_max_ can help avoid inappropriate initial determinations of the cell candidate. However, since a subsequent phase using a probabilistic model eliminates false positives, it is unnecessary to overly filter at this stage. A relatively broad range for *r*_min_ and *r*_max_ would suffice. In the final step of the initial candidate creation phase, for each obtained cell candidate or signal peak, its shape is carved out from the corresponding LoG filter using the watershed method, establishing the footprint of the cell candidate.

**Figure 4:**
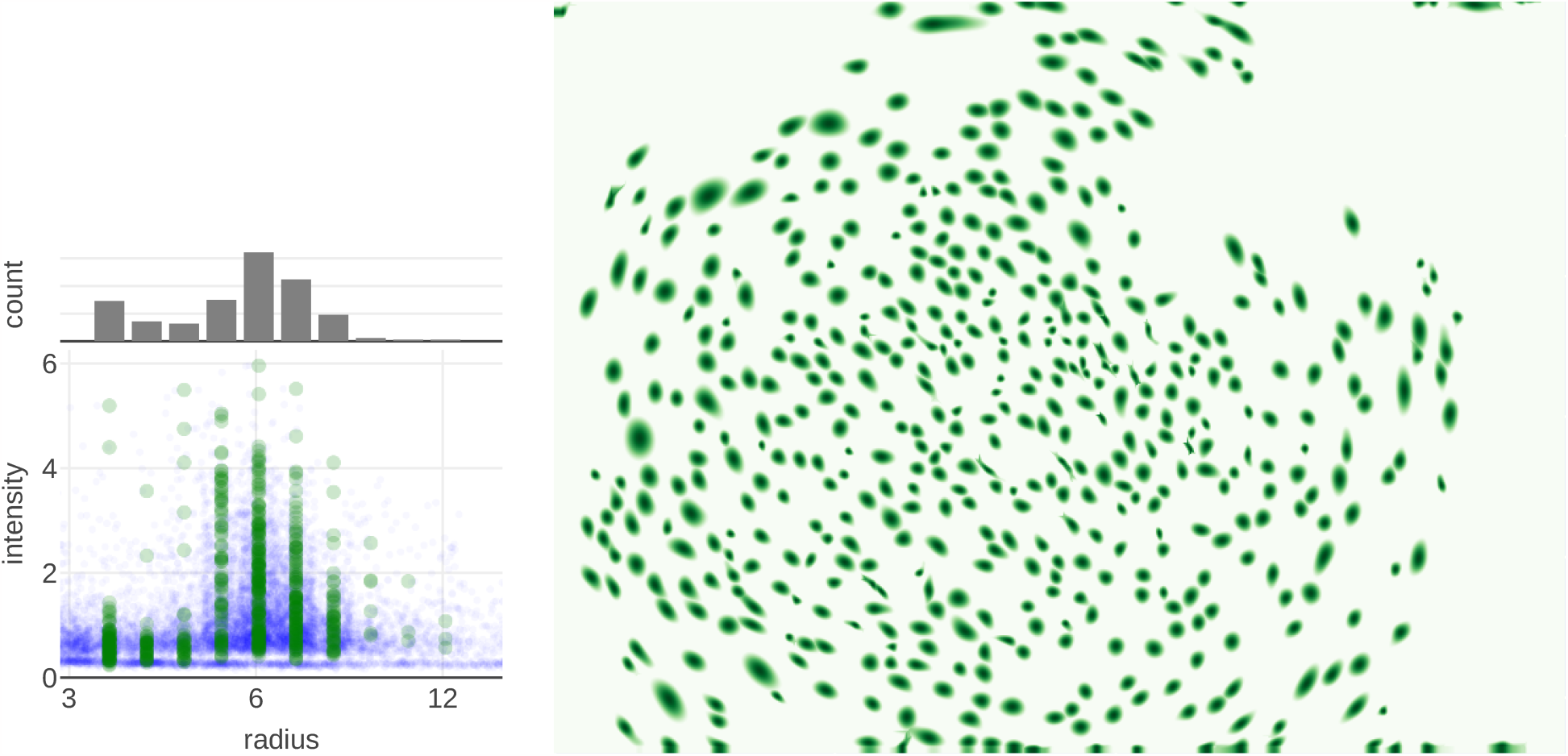
Initial footprints obtained by applying the proposed image processing process to Data 1. Left: The distribution of LoG filter radius *r* and intensity for each pixel’s obtained cell candidate (in blue) and cell candidates filtered by settings of *r*_min_, *r*_max_ (in green). Right: The footprints obtained for each cell candidate are overlaid and displayed.

Similarly, the initial step of creating cell candidates was applied to Data 2 (see Figure 5). Contrary to Data 1, there is an overlap between cells and other elements in the smaller region of *r*. This is believed to be due to the nature of two-photon microscopy, where elements such as cross-sections of axons are included in the recording. The probabilistic model-based optimization can clarify shapes and other features, making it somewhat possible to distinguish between minor non-cellular activities and actual cells. However, at present, there is a noticeable overlap based on radius information, and a clear distinction cannot be made regarding calcium activity. Whether to set *r*_min_ small to include all minor activities for analysis or set *r*_min_ large to focus only on activities that are definitely cellular varies depending on the research objective.

**Figure 5:**
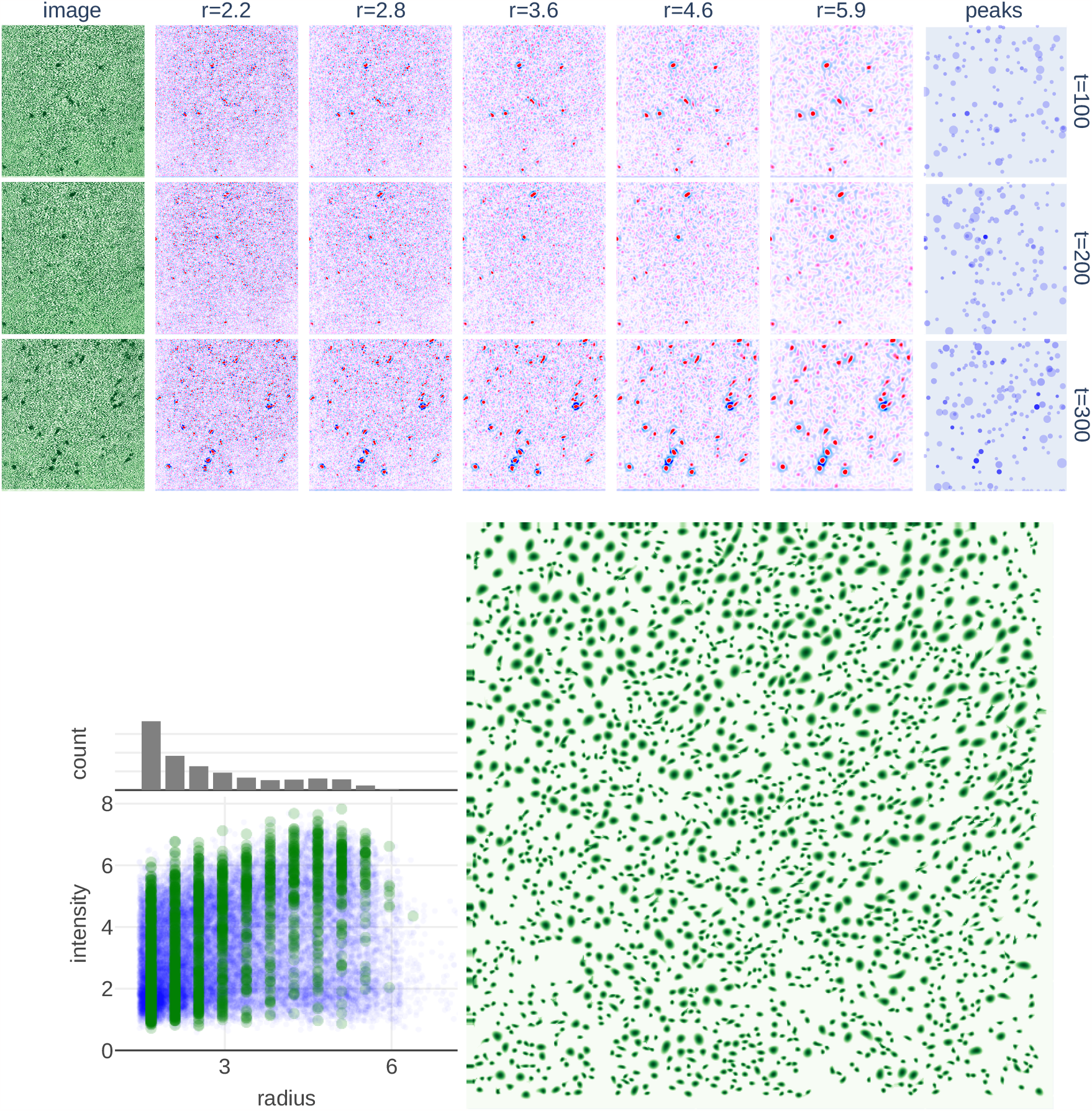
Initial footprints obtained by applying the proposed image processing process to Data 2 (correspoinding to the results adapted to Data 1 in Figure 3 and Figure 4). Top: For frames at t=100, 200, and 300, the original images and the results of applying LoG filters with radii r=2.2, 2.8, 3.6, 4.6, and 5.9 are displayed. The far-right column, labeled “peaks”, shows the detected maxima in a three-dimensional space with spatial dimensions (x, y) and radius r for each frame. The radius of the circle shown corresponds to r, and its transparency corresponds to intensity. Bottom-Left: The distribution of the LoG filter radius *r* and intensity for each pixel’s obtained cell candidate (in blue) and cell candidates filtered by settings of *r*_min_, *r*_max_ (in green). Bottom-Right: The footprints obtained for each cell candidate are overlaid and displayed. The detected cell candidates are divided into two groups: one with high intensity and a radius of 3 to 6 pixels, and the other with low radius or intensity. The former is expected to correspond to neurons and the latter to microstructures such as axons and other cross sections.

Additionally, the initial step of creating cell candidates was applied to Data 3, just as it was for Data 1. Numerous cell candidates, including their shapes, have been extracted from Data 2 (see Figure 6). However, the video includes local activity that changes over time, such as blood vessels, which are not neurons. At this initial candidate stage, many false negatives are expected to be included. In fact, compared to Data 1, the distribution of overlapping non-cellular elements can be seen on the larger side of the radius *r*. These imperfect results are expected to be refined during the optimization step using the probabilistic model.

**Figure 6:**
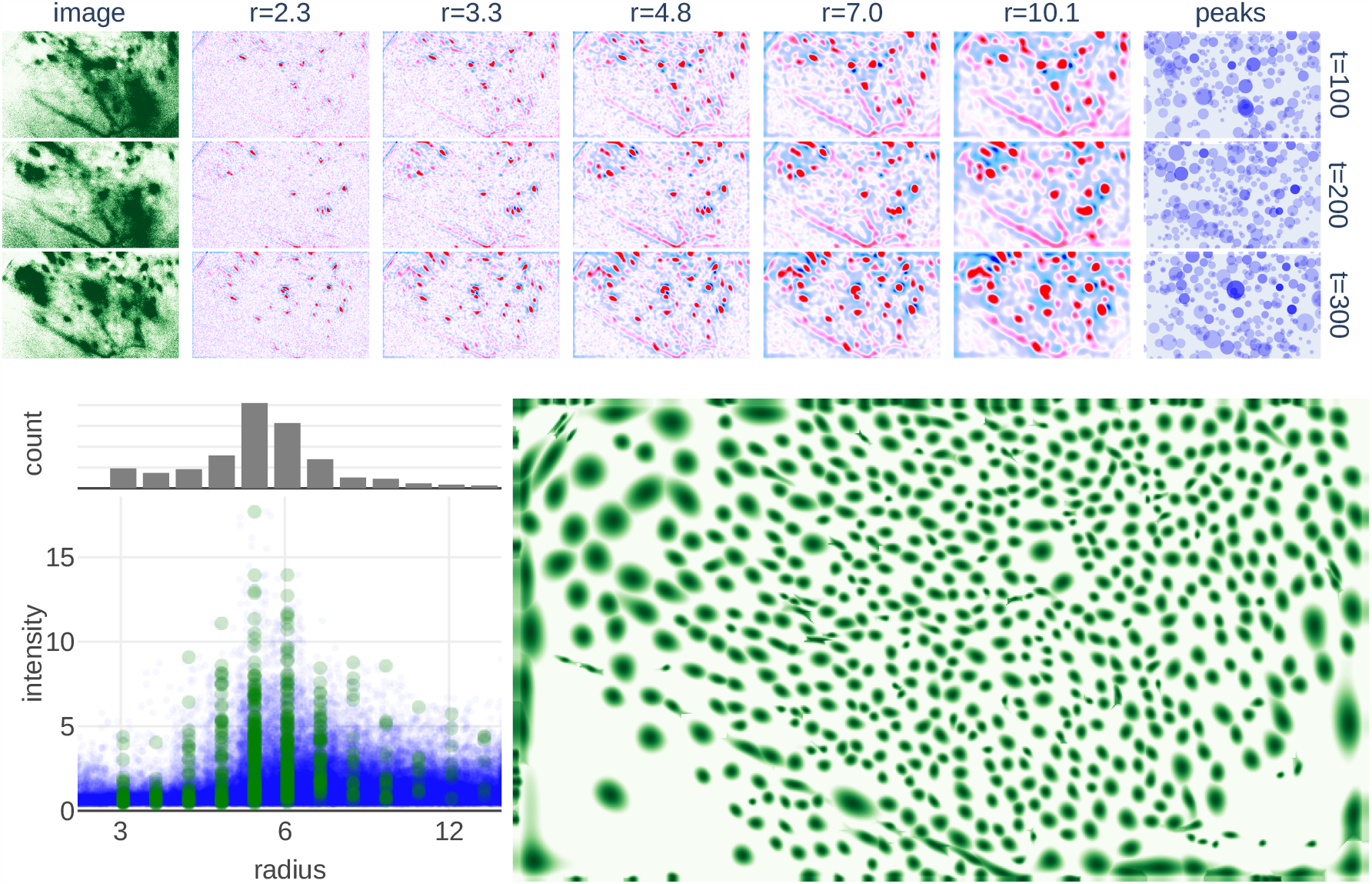
Initial footprints obtained by applying the proposed image processing process to Data 3 (corresponding to the results adapted to Data 1 in Figure 3 and Figure 4). Top: For frames at t=100, 200, and 300, the original images and the results of applying LoG filters with radii r=2.2, 3.3, 4.8, 7.0, and 10.1 are displayed. The far-right column, labeled “peaks”, shows the detected maxima in a three-dimensional space with spatial dimensions (x, y) and a radius r for each frame. The radius of the circle shown corresponds to *r*, and its transparency corresponds to intensity. Bottom-Left: The distribution of the LoG filter radius *r* and intensity for each pixel’s obtained cell candidate (in blue) and cell candidates filtered by settings of *r*_min_, *r*_max_ (in green). Bottom-Right: The footprints obtained for each cell candidate are overlaid and displayed. It can be seen that both the large-radius and elongated cell candidates are included. The former is likely to be an artifact caused by distortion in the peripheral field of view of the lens, while the latter is likely to correspond to activity such as blood vessels. These false-positive candidates should be removed from the final output.

### 4.2 HOTARU consistently eliminates false positives and improves cell shape

Applying the optimal temporal step to Data 1’s footprint, as acquired in the previous section, yields a spike time series (see Figure 7ABC). An evaluation of the signal intensity and spike density of the obtained spike time series reveals the inclusion of candidates with low signal strength and low signal-to-noise ratios, which are characterized by high density (refer to Figure 7D). Furthermore, an improved footprint is obtained when a spatial step is applied to the spike time series. Examination of the footprint’s “firmness” reveals that candidates with low signal intensity also possess low firmness. Regarding the radius, it is observed that the candidate cells predominantly fall within a reasonable range of 3-10 pixels.

**Figure 7:**
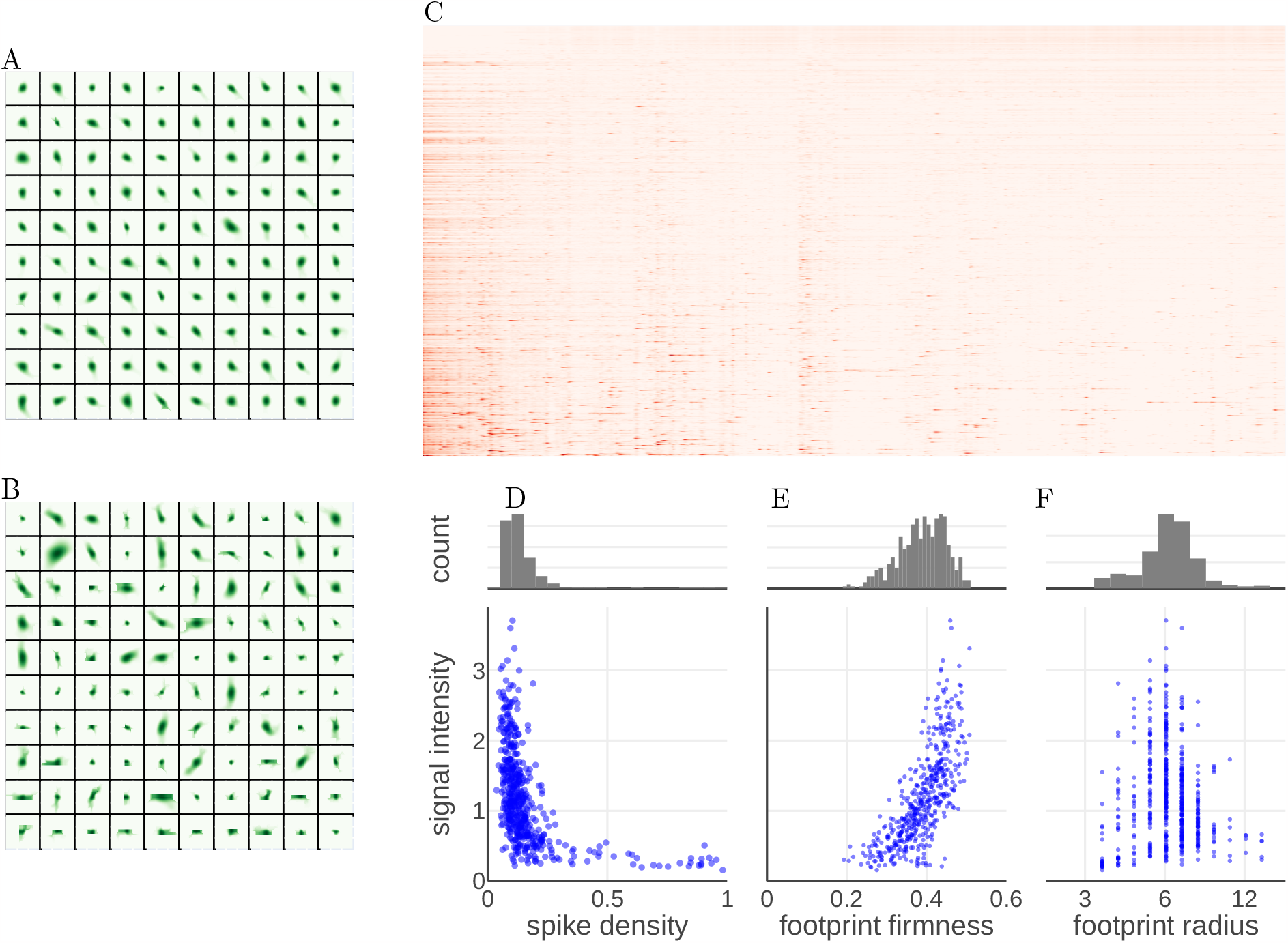
Results after one optimization step applied to initial cell candidates. A: Top 100 initial footprints with high intensity. B: Top 100 initial footprints with low intensity. C: Spike train estimated from the initial footprints. D: Distribution of spike density corresponding to C, shown as scatter plots and histograms combined with signal strength. EF: Distribution of firmness and radius estimated from C, shown as scatter plots and histograms combined with signal strength. At this stage, false-positive cell candidates with high density and low firmness are included. These candidates are removed through the optimization process without arbitrarily manipulating them.

The standard optimization parameters (*ζ*_*T*_ = *ζ*_*X*_ = 0.1, *λ*_*A*_ = 0.01, *λ*_*U*_ = 0.001, *λ*_*B*_ = 0.0001) were adopted. Setting *ζ*_*T*_ and *ζ*_*X*_ to 0 leads to significant numerical errors during iterations via the proximal operator, resulting in non-convergence or exceedingly slow convergence. On the other hand, setting *ζ*_*T*_ = *ζ*_*X*_ = 0.1 ensures a stable error reduction, which corresponds to regularization of the Hamiltonian eigenvalues by the baseline L2 regularization term, preventing them from reaching zero. No significant differences were observed when comparing the results between *ζ*_*T*_ = *ζ*_*X*_ = 0.1 and *ζ*_*T*_ = *ζ*_*X*_ = 0.2. Hence, all subsequent results are calculated with *ζ*_*T*_ = *ζ*_*X*_ = 0.1.

The results after 10 iterations of the temporal and spatial steps are presented in Figure 8. Following the spatial step, the cell regions are extracted, and the features are calculated using the LoG filter. Cell candidates with radii less than *r*_min_ are removed, while those with radii greater than *r*_max_ are treated as local background activity in subsequent steps. No cell removal, integration, or division, other than classification by minimum and maximum radii, has been arbitrarily performed. Consequently, candidates with high density or low firmness are automatically removed, resulting in unimodal distributions for the density, firmness, and radius metrics.

**Figure 8:**
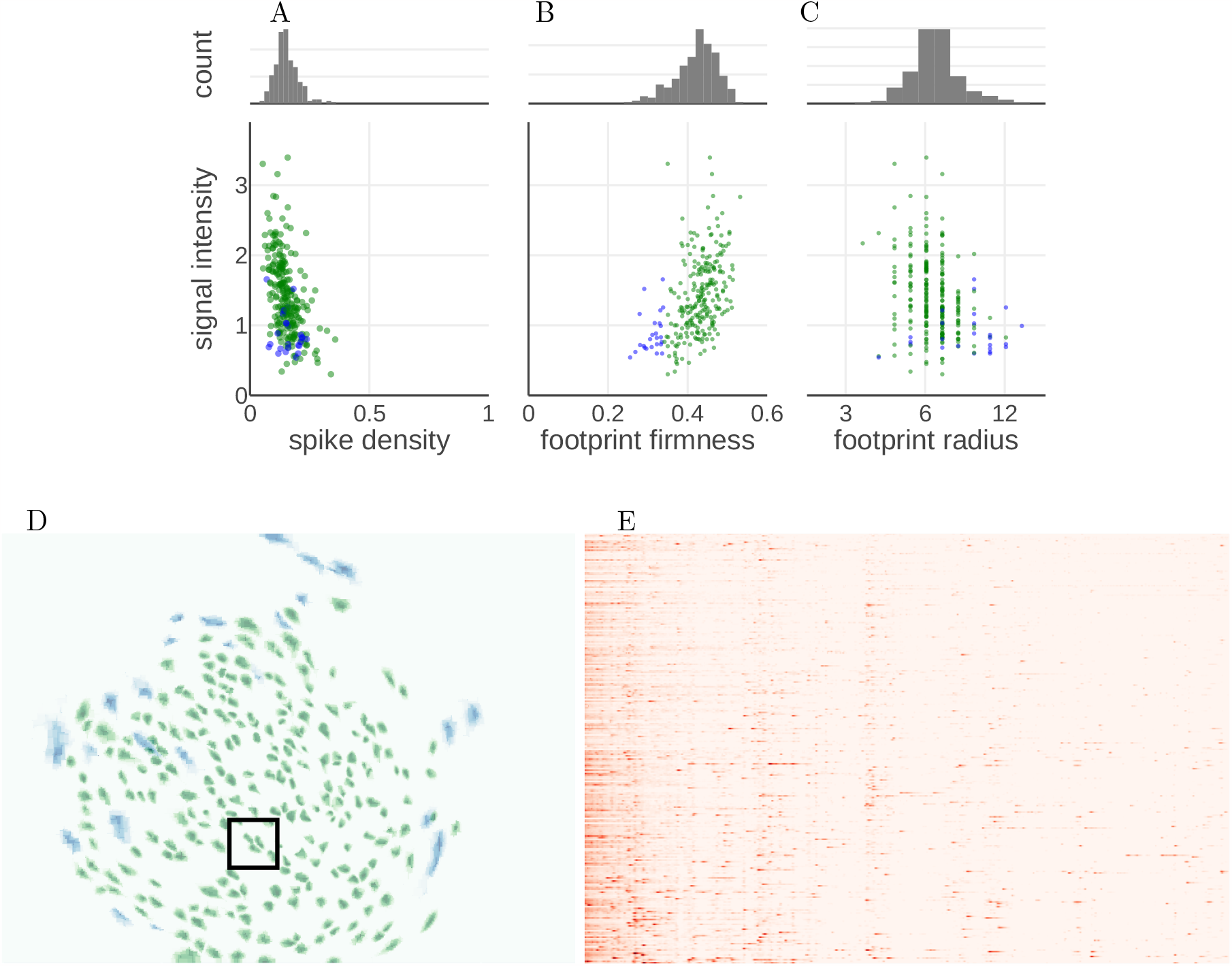
Final set of cell candidates. Compared to Figure 1, A, B, C: Distribution of spike density, footprint firmness, and footprint radius corresponding to Figure 1DEF, displayed as scatter plots and histograms combined with signal strength. Cell candidates with a firmness below 0.34 are highlighted in blue for reference. D: Overlay of candidate cell footprints, color-coded according to the points in the ABC scatter plots. A rectangular region surrounds the overlapping footprints of three cell candidates, whose details are shown in Figure 3. E: The resulting spike time series. Candidates with low firmness also exhibit low density and show distinct spike activities in their time series. These candidates are presumed to be cells distorted by lens aberrations. However, if the goal is to minimize false positives, the option of excluding these candidates from the analysis may be considered.

Signal intensities show substantial variability, and candidates with very low signal strength are also present. Generally, regularization introduces a penalty proportional to the signal strength, which could reduce the overall signal intensity or excessively lower the density of candidates with high signal intensity. However, candidates across a wide range of signal intensities exhibit similar density distributions, indicating that the group-normalized L1 regularization proposed in this paper is functional.

The initial number of candidate cells was 517, while the final count was 224. We evaluated the possibility of inadvertently subdividing the same cell based on criteria such as the distance between centers and the values of other footprints at the centers and found that although candidates likely to be subdivided were observed during iterations, they were automatically eliminated in the final results. With standard settings of CNMF-E, approximately 92 cells are identified; our proposed initial candidate detection and optimization process can be said to effectively detect neural cells with low signal intensity or low firing frequency with high precision.

### 4.3 Crosstalk elimination

When considering the cellular activity and corresponding calcium fluctuations simply as a weighted average with the footprint, crosstalk can occur in candidates where parts of the footprints overlap. In such cases, the activity of adjacent cells can be falsely detected even when a cell is not actually active, given that our proposed method based on CNMF is capable of eliminating such crosstalk. Figure 8D highlights a region encompassing three cells where this phenomenon can be observed. We compare the simple calcium fluctuations estimated by the weighted average of the footprints with the spike time series estimated by our method (see Figure 9). It becomes evident that the crosstalk observed in the calcium fluctuations is substantially reduced in the estimated spike time series. Although it is not completely eliminated, increasing *λ*_*U*_ can further reduce crosstalk; however, this comes with the trade-off of losing details in small fluctuations.

**Figure 9:**
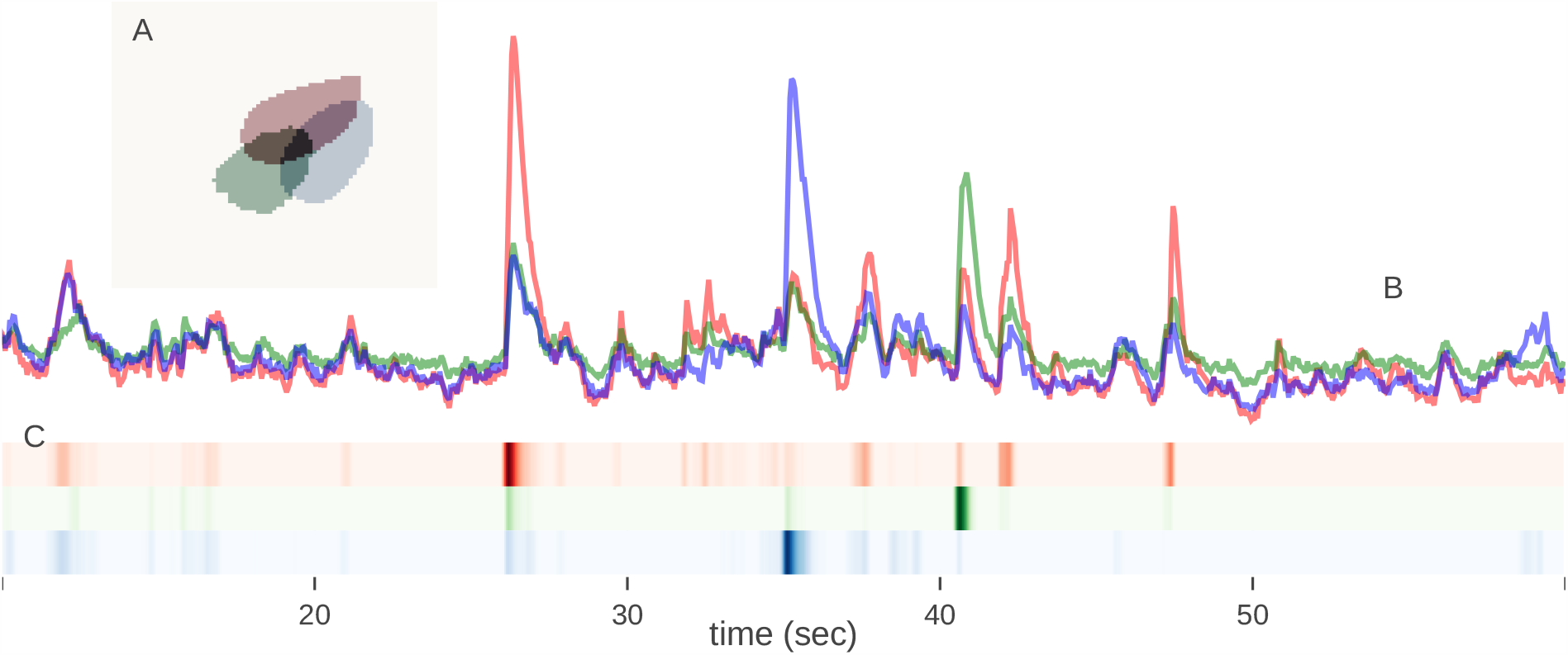
Calcium fluctuations and estimated spike sequences corresponding to cells with partially overlapping footprints. A: Footprint areas of the three cells. B: Weighted average of calcium fluctuations by footprint, where the colors correspond to the colors of the footprints (red, green, blue). C: Estimated spike time series, where the intensity of the color represents the spike intensity *u* at each time point.

### 4.4 Effects of regularization parameters and introduction of local background activity

To investigate the impact of the introduced regularization term, we examined the behavior during the optimization steps by varying the regularization parameter. We also compared cases in which cell candidates exceeding *r*_max_ were considered local background activity versus those in which they were removed. When *λ*_*A*_ was set to 0.0 or 0.001, the number of cell candidates did not converge despite repeated iterations, exhibiting a slight but continuous decrease. Figure 10 displays results up to 18 iterations; however, convergence was not achieved even after 80 iterations. This is attributed to the loss of firmness in the footprint during the spatial step. Increasing *λ*_*A*_ improves the firmness of actual neurons while making the shape of false positives unstable, thus accelerating the filtering of cell candidates.

**Figure 10:**
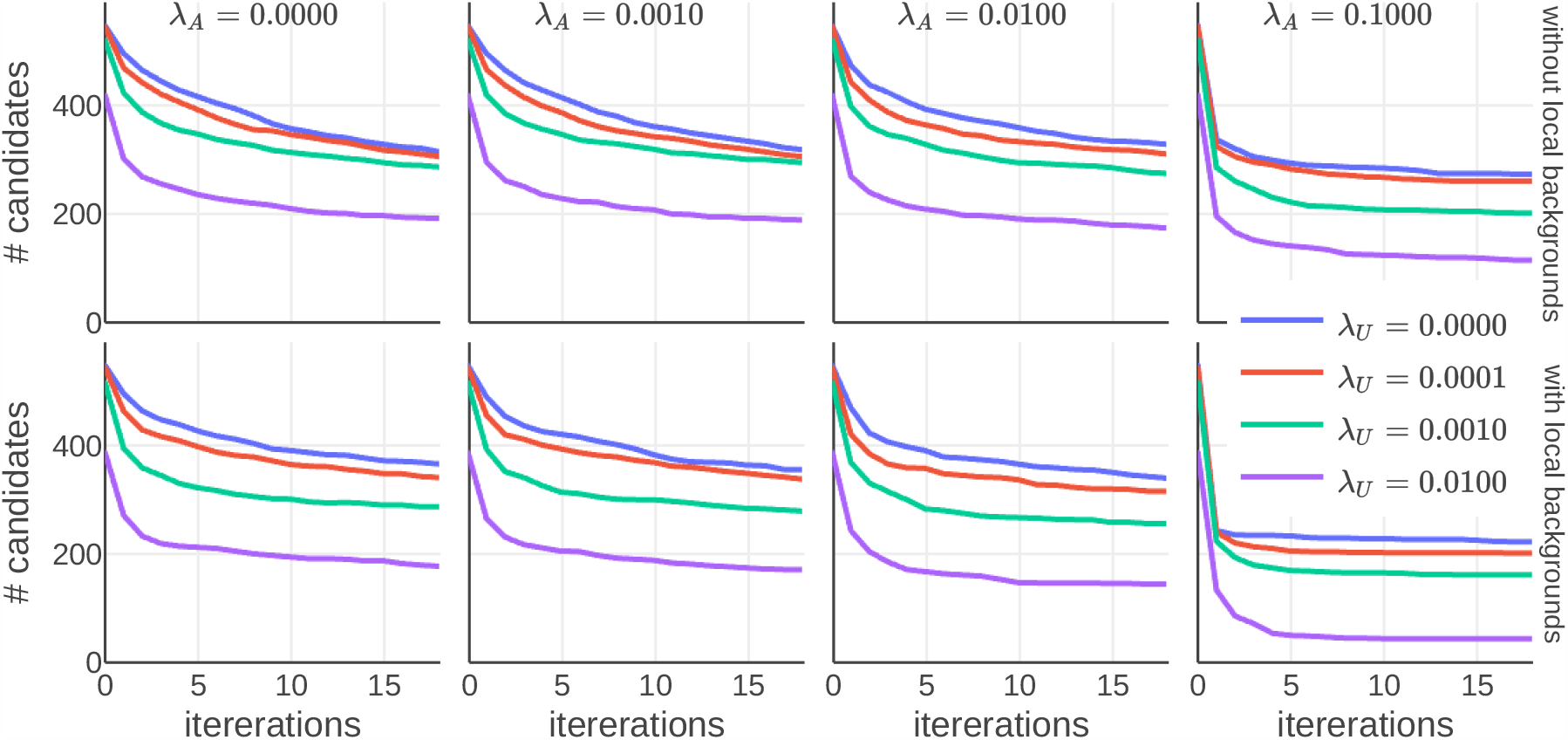
Changes in the number of cell candidates during iterations when varying *λ*_*A*_ and *λ*_*U*_ for Data 1. The top row represents cases where cell candidates exceeding *r*_max_ were removed, while the bottom row considers them as local background activity. The standard parameters for the entire study are set with local background activity at *λ*_*A*_ = 0.01 and *λ*_*U*_ = 0.01.

In this study, the introduction of local background activity at the standard parameter *λ*_*A*_ = 0.01 resulted in an easier convergence. *λ*_*U*_ has an effect similar to *λ*_*A*_ on the footprint, in that it suppresses the density of the spike time series. When *λ*_*U*_, the filtering of candidate cells is enhanced, and those with a low signal-to-noise ratio tend to be eliminated. However, even when the scale of *λ*_*U*_ was altered by 10 to 100 times, the estimate itself did not break down, indicating a trade-off between improving detection power and tolerating false positives.

Additionally, a trend was observed where larger regularization parameters resulted in shorter computation times within a single iteration. Coupled with the early narrowing down of cell candidates, the proposed algorithm also proves to be efficient in terms of computational time.

### 4.5 Handling of microstructures revealed by two-photon microscopy

Optimization step iterations were also applied to Data 2 obtained by two-photon microscopy (see Figure 11). Convergence was reached with fewer steps in Data 2 compared to Data 1, which is expected to correspond to a lesser extent of overlapping cells. Additionally, no local background activity was observed in Data 2. Variations in *λ*_*A*_ and *λ*_*U*_ resulted in less change compared to Data 1; in the range of 0 to 0.01 for both *λ*_*A*_ and *λ*_*U*_, the number of detected cells did not fluctuate significantly. This is likely due to the distinct characteristic of two-photon microscopy that provides a clear focus, making it easier to discern false positives. However, at *λ*_*A*_ = 0.1, the number of detected cells drastically decreased. This is because the cells in Data 2 are smaller in shape compared to Data 1, making the effect of *λ*_*A*_ relatively stronger. Similarly, factors such as sampling frequency could also influence the effects of *λ*_*U*_. Although tuning optimal *λ*_*A*_ values from parameters such as sampling frequency or initial cell candidate radius could be possible, this paper emphasizes the system’s stability across a broad parameter range and does not delve into tuning specifics.

**Figure 11:**
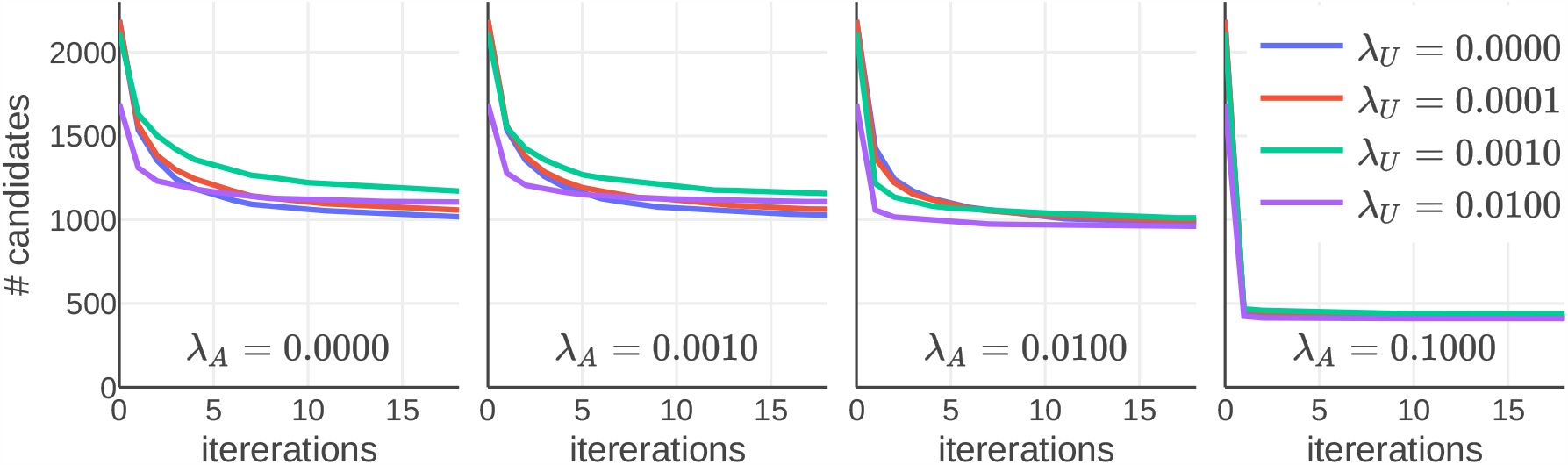
Changes in the number of cell candidates during iterations when varying *λ*_*A*_ and *λ*_*U*_ for Data 2. Although local background activity was considered, none was actually detected.

In Data 2, optimization iterations led to the removal of initial cell candidates with high spike density and low footprint firmness. The remaining cell candidates exhibited improved intensity and firmness while showing decreased density. However, unlike data 1, where only neurons were automatically detected through optimization iterations, data 2 still contained small microstructures post-optimization. Using Gaussian mixture models for clustering based on features such as firmness, density, and intensity roughly separated the groups into neurons and microstructures (see Figure 12ABC). Of the 1,017 candidates obtained through optimization, 460 were clustered as neurons. Visual inspection of the footprints also suggested reasonable discrimination between cells (green) and local structures (blue) (see Figure 12D). For a more precise classification, manual labeling of neurons and microstructures followed by the generation of machine learning models could be implemented.

**Figure 12:**
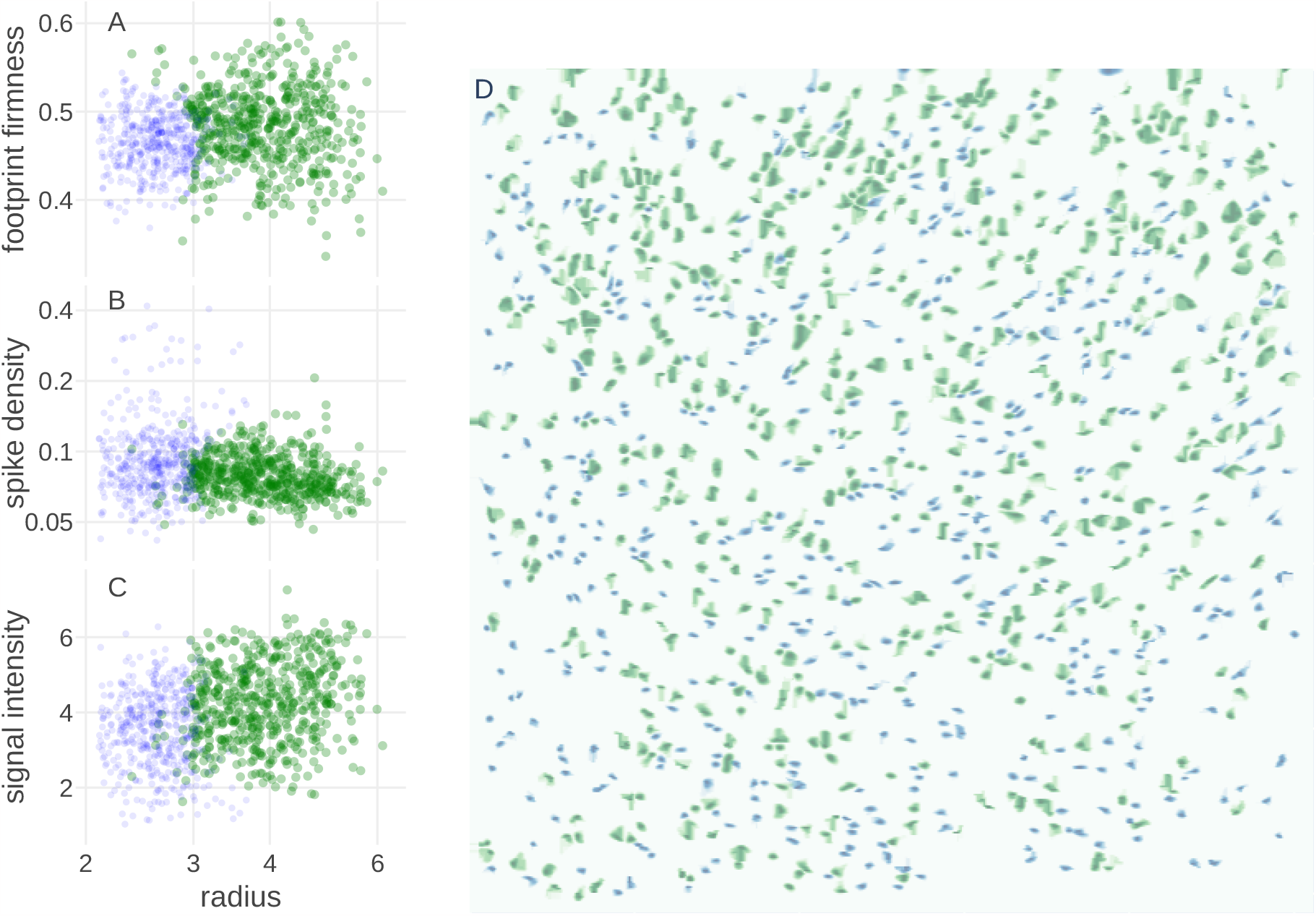
Optimization and microstructure separation for Data 2. ABC: Distribution of radius, firmness, density, and intensity for cell candidates obtained after optimization iterations. Notably, the distribution mainly separates into two groups based on radius. The green and blue results are from Gaussian mixture models clustering. D: Overlaid footprints of cell candidates. The green and blue footprints correspond to individual points in ABC.

### 4.6 Mitigating the influence of vascular on neuronal identification

Continuing the analysis, iterative optimization steps were applied to Data 3. In Data 3, the vascular movements are prominently visible in the videos. When local background activity was not considered and cell candidates with a radius greater than *r*_max_ were eliminated, the estimation of neurons was significantly compromised, resulting in a continuous reduction of cell candidates (see Figure 13). Furthermore, when standard parameters were applied for the optimization steps, several instances were observed where structures visually similar to blood vessels were detected as cell candidates (see Figure 14A).

**Figure 13:**
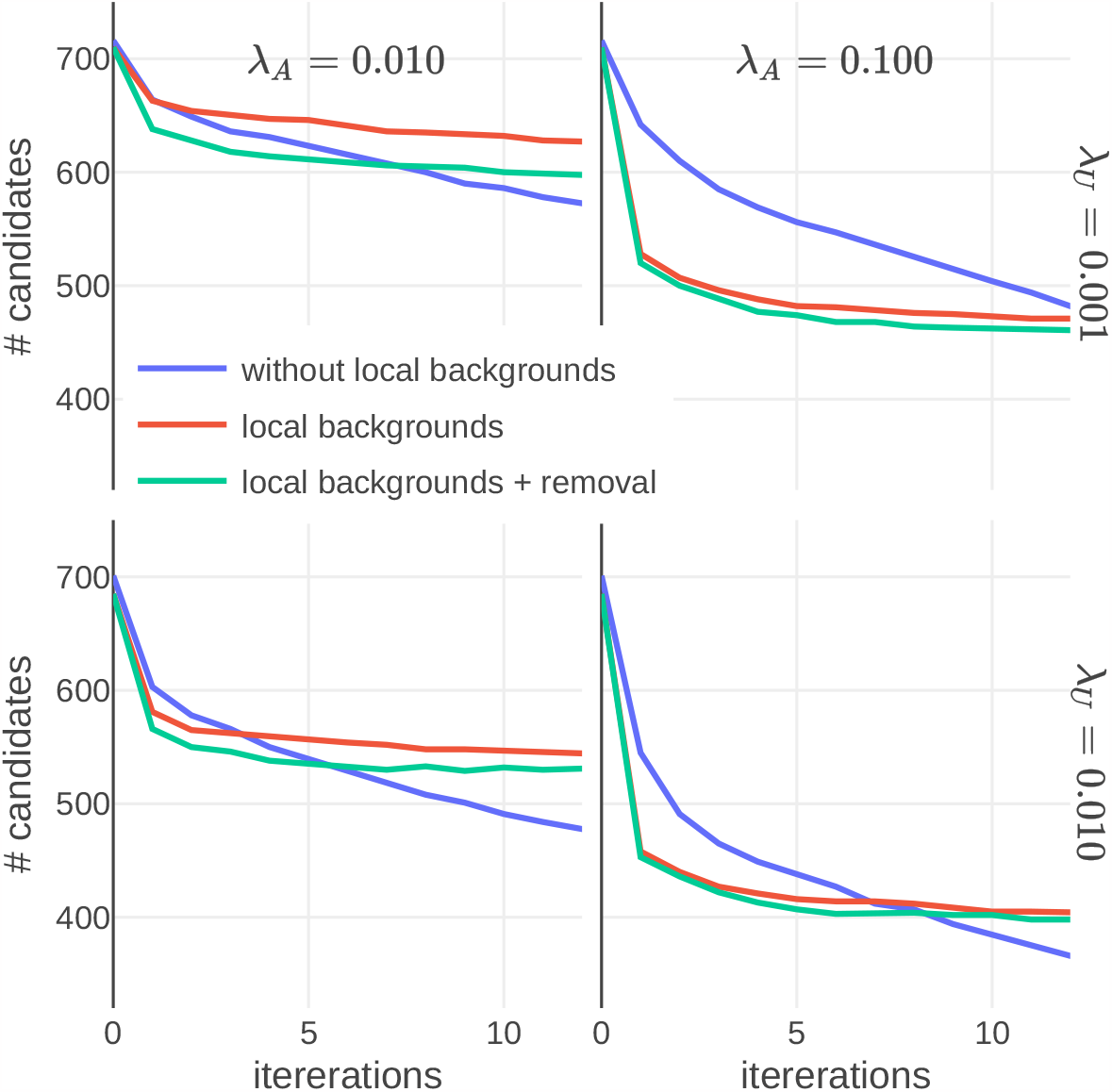
Variation in the number of cell candidates over iterations when changing the regularization parameters *λ*_*A*_ and *λ*_*U*_ in Data 3. The number of cell candidates decreases when candidates with a radius greater than *r*_max_ are removed (blue), as opposed to when they are considered local background activity (red). Additional aggressive measures to classify some cell candidates as local background activity are indicated in green. Without considering local background activity, the number of cell candidates does not converge. With smaller regularization parameters, more cell candidates are classified as local background activity, but the difference decreases with larger regularization parameters.

**Figure 14:**
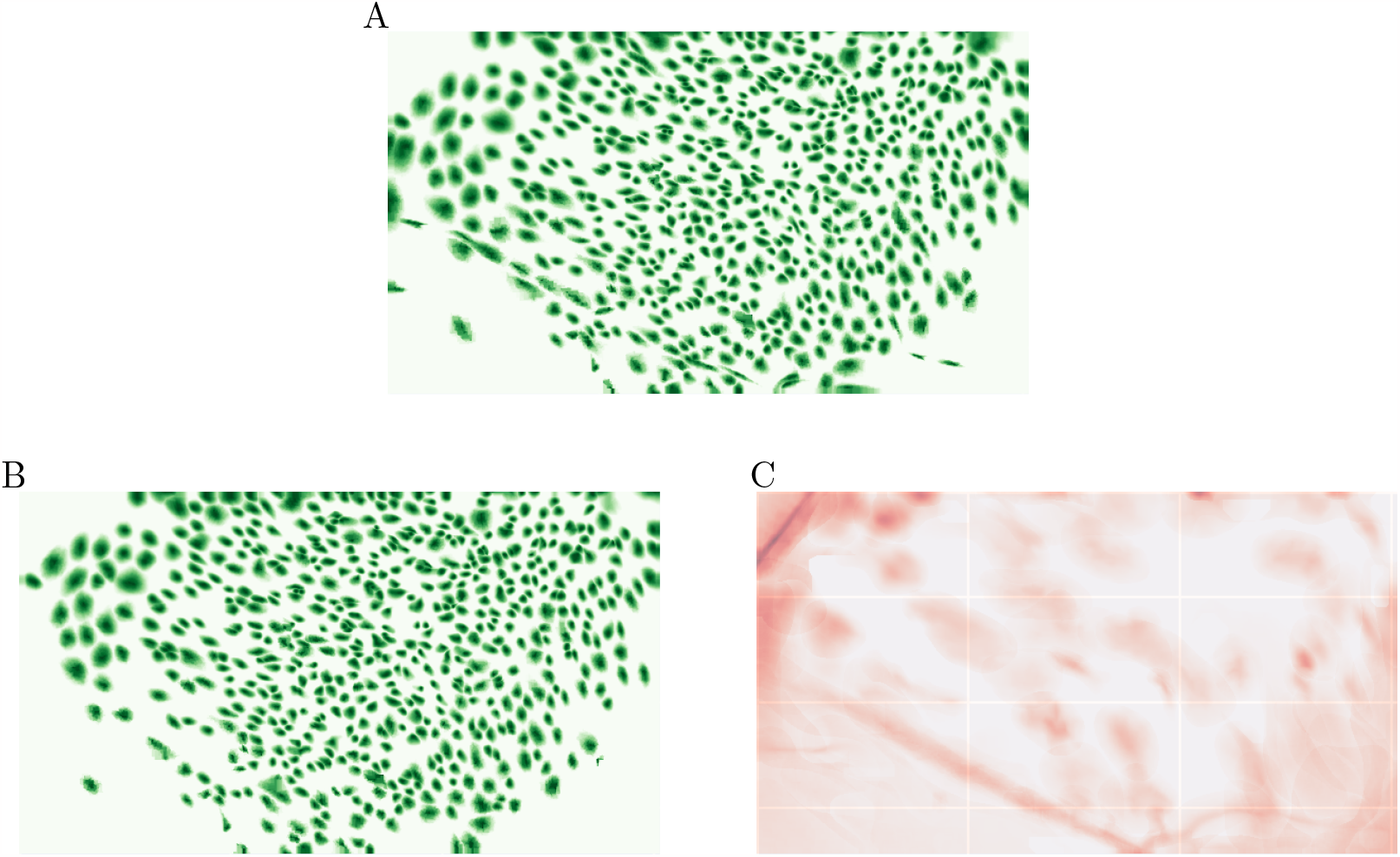
Footprints and local background activity in cell candidates for Data 3. A: Footprints obtained by applying the optimization steps in the same manner as in Data 1 and Data 2. B: Footprints of cell candidates classified as neurons or local background activity according to their elliptical shape. C: Corresponding local background activity to B.

To address these issues, additional measures were employed. Specifically, the footprints were approximated as ellipses, and the cell candidates were considered to have local background activity if the ratio of their major to minor axes was three times greater or greater. Furthermore, the distribution of “firmness” and “density” was approximated as a multivariate normal distribution, and cell candidates further than 4*σ* from the center of the distribution were also considered local background activity. Approximately 30 candidates met this criterion in ten iterations. The 600 resulting cell candidates showed distinct cellular morphology and spiking activities ((see Figure 14B)), and vascular activities were clearly delineated as local background activities (see Figure 14C). Indirect evidence for the validity of these additional measures was observed when increasing the regularization parameters *λ*_*A*_ and *λ*_*U*_, which resulted in fewer cell candidates being classified as local background activity (the difference between red and green in Figure 13).

### 4.7 Computational Time Analysis for Large-Scale Dataset

In the case of Data 2, the most sizable among the datasets examined, we carefully measured the computational time involved for both the initialization and the subsequent 10 optimization steps. The total computational duration was clocked at 2517 s. The individual components of this time expenditure were further broken down for analysis. Specifically, data pre-processing and the generation of the initial footprint accounted for 250 s. The first temporal step consumed a total of 205 s, with a notable 68 s allocated for preliminary matrix calculations involved in the optimization problem. Additionally, 60 s were required to complete a full scan of the 29 GB dataset stored on disk. The first spatial step required 232 s and the matrix calculations consistently consumed 68 s, as observed in the temporal step. It was observed that in subsequent iterations, the computational time exhibited a decreasing trend, which could be attributed to factors such as the reduction in cell count. This detailed temporal breakdown serves not only as an assessment of the current algorithm’s efficiency but also provides valuable guidelines for computational optimizations in future work. HOTARU was run on an Intel Xeon W-2123 at 3.60 GHz with 128 GB memory and Ubuntu 20.04.2 LTS on NVIDIA TITAN RTX.

HOTARU is implemented in Python and “jax” and makes effective use of GPUs. The implementation is released as open-source software (https://github.com/tk2lab/HOTARU), ver. 5, which corresponds to that used in this study.

## 5 Discussion

In this study, we present the HOTARU system. This novel approach to identifies candidate cells in video frames by specifying a set of candidate radii 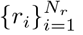, minimum radius *r*_min_, maximum radius *r*_*max*_, and relative intercellular distance *θ*. This system enables the detection of numerous cell candidates without overlap, effectively narrowing down to a realistic number of cells. The validity of these parameters can be easily verified by calculating the radius and intensity distribution of the cellular shapes. Traditional cell candidate detection algorithms struggle with setting an appropriate threshold for candidate adoption, as lowering the threshold to include more candidates often leads to a dramatic increase in the number of false positives. Our approach avoids this problem by allowing the fine-tuning of *θ* based on whether the focus is on detecting as many cells as possible or minimizing false positives. Furthermore, subsequent optimization steps automatically remove false positives, reducing the need for meticulous tuning.

In the optimization steps, we stabilized the solution by carefully setting temporal and spatial baselines for the entire video. We introduced a new normalization method tailored to each cell candidate. When combined with a corresponding proximal operator, this method enhances the estimation of low-SNR cell candidates without sacrificing the accuracy of high-SNR cells. We also integrated the optimization of local background activity, enabling the detection of neurons even in the vicinity of vascular structures. One significant advantage of our approach is its consistent convergence within roughly ten iterations across all datasets. This feature reduces sensitivity to local solutions and parameter settings – an essential aspect when analyzing calcium imaging data where ground truth may be unavailable.

In addition, we propose evaluation metrics such as signal intensity, radius of the footprint, firmness of the footprint, and spike density alongside our analytical method. The unimodal distribution of these metrics without arbitrary manipulation attests to the validity of our approach. Variations in the regularization parameters for cellular shape and spike time series correlate with the number of low-SNR cell candidates included, offering a trade-off that can be adjusted based on the specific aims of the experiment. Although minimizing false positives is generally preferable, the ability to tailor parameters according to objectives is often advantageous. Even for cells with low signal intensity, the chances of false positives are not particularly high since their firmness and density remain within a certain range.

While this study primarily showcases a basic yet effective framework, there is a potential for further refinements. Detailed parameter tuning and improvements in identifying microstructures or vascular features in Data 2 and Data 3 could be addressed through the integration of more specialized, high-accuracy techniques.

## Acknowledgments

This work was supported by JSPS KAKENHI (grant number: JP18H05213), the Core Research for Evolutional Science and Technology (CREST) program (JPMJCR13W1) of the Japan Science and Technology Agency (JST), a Grant-in-Aid for Scientific Research on Innovative Areas “Memory dynamism” (JP25115002) from MEXT, and the Takeda Science Foundation’s support for K.I. T.F. work was supported by the Brain/Minds project from the MEXT, KAKENHI (JP15H04265) from the JSPS, a Grant-in-Aid for Scientific Research on Innovative Areas “Memory dynamics” (JP16H01289) and “Artificial Intelligence and Brain Science” (JP17H06036) from the MEXT. Y.H. work was supported by RIKEN, the Human Frontier Science Program (RGP0022/2013), the Core Research for Evolutional Science and Technology (CREST) program (JPMJCR20E4) of the Japan Science and Technology Agency (JST), a Grant-in-Aid for Scientific Research (S) (JP22H04981), a Grant-in-Aid for Scientific Research on Innovative Areas “Memory dynamism” (JP16H01292), “Synapse Pathology” (JP22110006), and “Multiscale Brain” (JP18H05434) from the MEXT. T.T. work was supported by KAKENHI (JP26870577, JP19K12104) from the JSPS.

## A Methods Details

### Algorithm 1 Find initial peaks

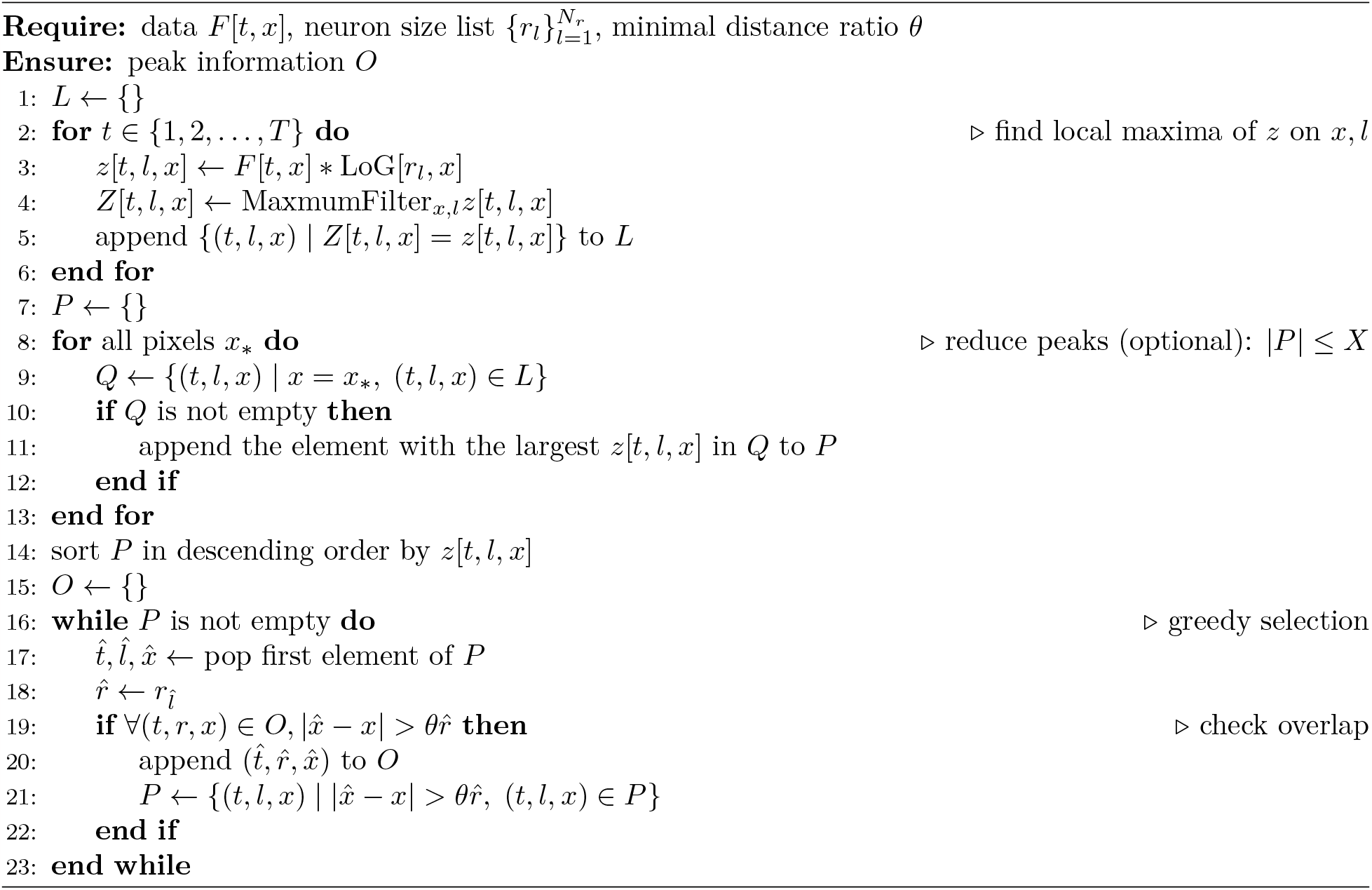

### Algorithm 2 Make intial footprints

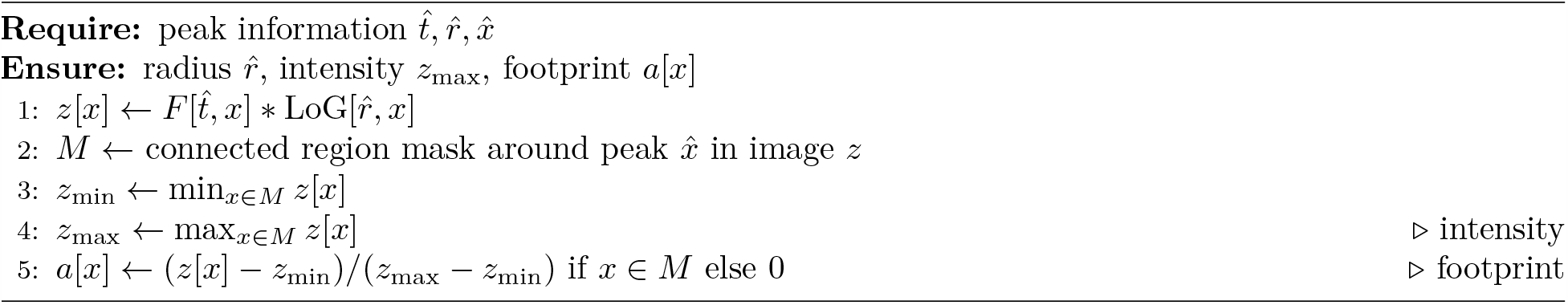

### Algorithm 3 Clean up footprints

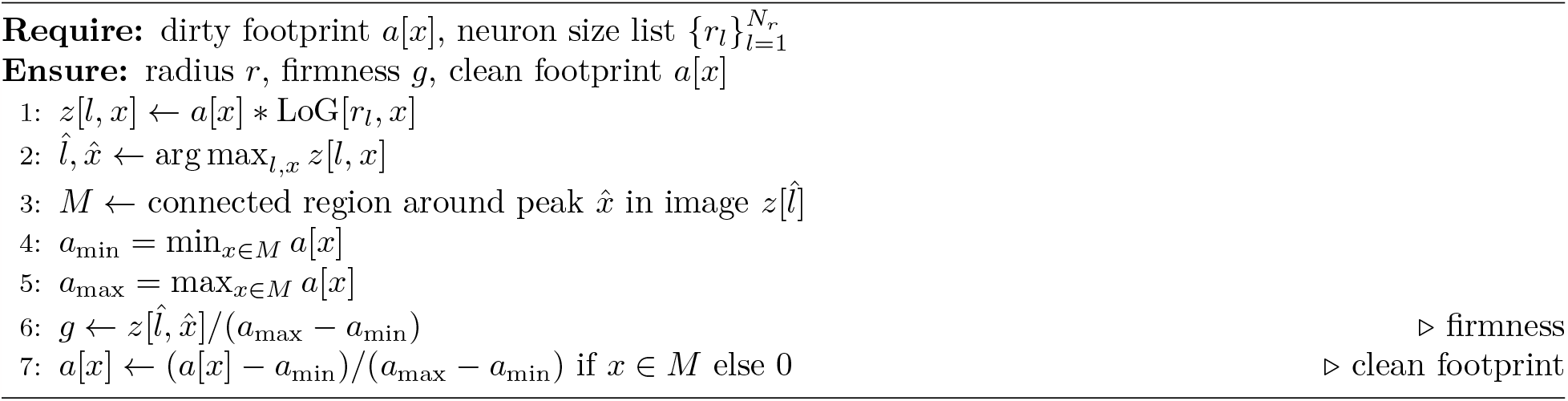

### A.1 Finding cell candidates from video data using LoG filters

To identify the initial cell candidates, we began by preparing a list of cell size parameters 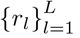. We applied the LoG filter corresponding to all *r*_*l*_ to the image *f*_*t*_[*x*] at time *t* resulting in *L* types of filtered images. This collection of images is considered to be a 3D structure of *L× W× H*, and all maxima and their positions in the 3D space are collected.

Since the peak information collected is huge, it is efficiently condensed. First, the collected peaks are grouped by position *x* in the image, and only those with the largest maxima are retained. Currently, the cell size *r* of the adopted peak is also recorded. Subsequently, the remaining peaks are sorted and adopted in descending order by the maximum value, which corresponds to signal intensity. However, if the distance between the centers is within *θ* times the radius of the peaks with stronger signals than those already selected, the peaks are not selected. The detailed procedure is shown in Algorithm 1.

Following this, we determine the specific cell shape based on the obtained peak position and filter size. Once again, an LoG filter corresponding to the filter size is applied, and the watershed cut algorithm is applied to the center of the peak position to extract the cell region. The filter image is cropped in the extracted region and normalized to [0, 1] as the initial footprint. These details are shown in Algorithm 2.

The iterative footprint is also improved using a LoG filter and the watershed cut algorithm. Again, multiple LoG filters are applied to each footprint. The maximum peak is then found, and the watershed cut algorithm is applied to the filtered image around the peak to extract the cellular region. The cell region is extracted from the original footprint and normalized to [0, 1] to obtain the improved footprint. These details are shown in Algorithm 3.

### A.2 Overall framework

The overall algorithm for HOTARU can be summarized as follows: The process begins with the identification of cell candidates (FIND) and the generation of initial footprints (INIT). Following this, temporal optimization steps (T-P) are executed. Subsequently, the algorithm undergoes a loop consisting of a spatial optimization step (S-P), footprint cleaning (CLEAN), and another temporal optimization step (T-P) as one complete stage. During the CLEAN process, cell candidates with a radius less than *r*_min_ are discarded, while those with a radius greater than *r*_max_ are considered local background activities.

In this study, the set of candidate radii 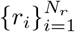 for Data 1 and Data 3 comprised 16 logarithmically equidistant values ranging from 2 to 16, specifically 2.00, 2.30, 2.64, 3.03, 3.48, 4.00, 4.59, 5.28, 6.06, 6.96, 8.00, 9.19, 10.56, 12.13, 13.93, and 16.00. For Data 2, which features smaller cells when translated to pixel units, 15 values between 2 and 8 were used, specifically 2.00, 2.21, 2.44, 2.69, 2.97, 3.28, 3.62, 4.00, 4.42, 4.88, 5.38, 5.94, 6.56, 7.25, and 8.00. The actual ranges for *r*_min_ and *r*_max_ were determined based on the distribution of the initial radii and intensities of the footprint. Specifically, *r*_min_ = 3.1, *r*_max_ = 15.9 for Data 1; *r*_min_ = 2.7, *r*_max_ = 15.9 for Data 2; and *r*_min_ = 2.1, *r*_max_ = 7.9 for Data 3.

For the optimization parameters, unless explicitly validated for their effects, the values *λ*_*A*_ = 0.01, *λ*_*U*_ = 0.001, *λ*_*B*_ = 0.0001, *ζ*_*T*_ = *ζ*_*X*_ = 0.1 were used in all datasets. During the optimization steps, the Nesterov accelerated proximal gradient method was used, and 100 steps of computation were considered to be one epoch. The iteration process ended when the rate of change in the objective function *E* was less than 10^−4^. Step sizes *η* were set to 10^−6^ *× T* for spatial steps and 10^−7^ *× X* for temporal steps. In general, each stage for all datasets required approximately 10-30 epochs to complete. As demonstrated in this manuscript, under standard parameters, cellular candidate numbers, shapes, and spike trains generally converge within approximately 10 stages.

### A.3 Complexity analysis

Although calcium imaging is now being recorded on a larger scale, it is safe to assume that a certain upper limit is set for spatial resolution and that the computational complexity in the time direction *T* is most important. Considering the computational complexity of a time-step update, the precomputation of the matrix is dominated by *O*(*KTX*) due to the matrix product of *F* and *A*. When applying the proximal gradient, the computation of *A* dominates *O*(*K*^2^*T*). For the space step, the pre-computation and the computational complexity per iteration, respectively, are *O*(*KTX*) and *O*(*K*^2^*X*). Since the number of cells *K* is proportional to *X* but independent of *T*, the computational complexity for *T* is *O*(*T*). For each frame, the process of finding the initial footprint can also be performed independently or in parallel, and using the shrinkage per pixel, subsequent operations are independent of *T*.

## B Materials Details

All procedures involving the use of animals complied with the guidelines of the National Institutes of Health, and they were approved by the Animal Care and Use Committee of the University of Toyama, the Institutional Committee for the Care, the RIKEN Animal Experiments Committee, and the Genetic Recombinant Experiment Safety Committee. Thy1-G-CaMP7-T2A-DsRed2 mice that express the fluorescent calcium indicator protein G-CaMP7 and the red fluorescent protein DsRed2 under the control of the neuron-specific Thy1 promoter were used. Details on the generation and characterization of the transgenic mice are described in Sato et al. (2020).

### B.1 Hippocampal data by miniature microscope

The mice were kept on a 12-h/12-h light-dark cycle (lights on 7:00 am) at 24*±*3^*°*^ C and 55 *±* 5% humidity, provided with food and water ad libitum, and housed with littermates until 1–-5 days before surgery. We performed hippocampal surgery for gradient refractive index (GRIN) lens setting as previously described (Barretto et al., 2011; Ghosh et al., 2011; Ziv et al., 2013; Ghandour et al., 2019). All surgery was conducted on approximately 12-week-old male Thy1::G-CaMP7-p2A-DsRed mice on a C57BL/6J background. The mice were anesthetized with a pentobarbital solution (80 mg/kg of body weight; intraperitoneal injection), and the fully anesthetized mice were placed in a stereotactic apparatus (Narishige, Japan). To set the cannula lens sleeve (outer diameter, 1.8 mm; length, 3.6 mm; Inscopix, CA), a craniotomy was performed with a diameter of 2.0 mm. The cylindrical column of the neocortex and corpus callosum above the alveus covering the dorsal hippocampus was aspirated using a 27-gauge blunt drawing up needle with saline. The cannula lens sleeve was gently placed on the alveus and fixed to the edge of the burr hole with bone wax, which was melted using a low-temperature cautery. The cannula lens sleeve targeted the right hemisphere (AP 2.0 mm, ML 1.5 mm at center). After setting the anchor screws onto the skull, we covered the skull with dental cement, which fixed the cannula lens sleeve to the skull and anchor screws.

Approximately 3–4 weeks after surgery, mice were anesthetized with isoflurane (1.5–2%), and a GRIN lens (outer diameter, 1.0 mm; length, 4.0 mm; Inscopix) was inserted into the cannula lens sleeve and fixed with ultraviolet-curing adhesive (Norland, NOA 81). The integrated miniature microscope (nVista HD, Inscopix) (Ghosh et al., 2011) with a microscope baseplate (Inscopix) was placed above the GRIN lens, at which G-CaMP7 fluorescence was observed. The microscope baseplate was fixed with the head of the anchor screw using dental cement. This shaded the GRIN lens, after which the integrated microscope was detached from the baseplate. The GRIN lens was covered by attaching the microscope baseplate cover to the base plate until calcium imaging was performed.

Calcium imaging was performed during the light cycle, and calcium events were captured at 20 Hz using nVista acquisition software (Inscopix). The integrated microscope was re-attached to the baseplate, and then the mice were introduced into a novel context consisting of a cylindrical chamber (diameter*×* height: 180 *×*230 mm^2^) with a white acrylic floor and walls covered with black tape. The captured video was processed as previously described (Kitamura et al., 2015; Ghandour et al., 2019). Motion correction by Mosaic software (Inscopix) was applied to the dF/F signal video.

### B.2 Two-photon calcium imaging

The two-photon microscopy images (Data 3) were acquired using a Nikon A1MP (Nikon, Tokyo, Japan) equipped with a 16x, NA 0.8 water immersion objective lens as described previously (Sato et al., 2020). Briefly, an optical window was surgically implanted in the cortex overlaying the dorsal hippocampus. G-CaMP7 and DsRed2 were excited at 910 nm using a Ti-sapphire laser (MaiTai DeepSee eHP, Spectra-Physics, Milpitas, CA). G-CaMP7 fluorescence was separated using a dichroic mirror at 560 nm and collected using an external GaAsP photomultiplier tube (10770PB-40, Hamamatsu Photonics, Japan). The microscope focused at approximately 150 μm from the hippocampal surface to image CA1 pyramidal cells labeled with G-CaMP7. A resonant-galvo scanner attached to the microscope acquired 512 *×* 12-pixel images at 15 frames per second. An imaging session lasted 10 minutes. The field of view was 532 *×* 532 μm.

